# Tuft cell-derived acetylcholine regulates epithelial fluid secretion

**DOI:** 10.1101/2023.03.17.533208

**Authors:** Tyler E. Billipp, Connie Fung, Lily M. Webeck, Derek B. Sargent, Matthew B. Gologorsky, Margaret M. McDaniel, Darshan N. Kasal, John W. McGinty, Kaitlyn A. Barrow, Lucille M. Rich, Alessio Barilli, Mark Sabat, Jason S. Debley, Richard Myers, Michael R. Howitt, Jakob von Moltke

## Abstract

Tuft cells are solitary chemosensory epithelial cells that can sense lumenal stimuli at mucosal barriers and secrete effector molecules to regulate the physiology and immune state of their surrounding tissue. In the small intestine, tuft cells detect parasitic worms (helminths) and microbe-derived succinate, and signal to immune cells to trigger a Type 2 immune response that leads to extensive epithelial remodeling spanning several days. Acetylcholine (ACh) from airway tuft cells has been shown to stimulate acute changes in breathing and mucocilliary clearance, but its function in the intestine is unknown. Here we show that tuft cell chemosensing in the intestine leads to release of ACh, but that this does not contribute to immune cell activation or associated tissue remodeling. Instead, tuft cell-derived ACh triggers immediate fluid secretion from neighboring epithelial cells into the intestinal lumen. This tuft cell-regulated fluid secretion is amplified during Type 2 inflammation, and helminth clearance is delayed in mice lacking tuft cell ACh. The coupling of the chemosensory function of tuft cells with fluid secretion creates an epithelium-intrinsic response unit that effects a physiological change within seconds of activation. This response mechanism is shared by tuft cells across tissues, and serves to regulate the epithelial secretion that is both a hallmark of Type 2 immunity and an essential component of homeostatic maintenance at mucosal barriers.

## Introduction

The physiologic function and immune defense of mucosal tissues require fluid secretion, and epithelial cells employ multiple independent mechanisms to regulate this process. For example, cyclic AMP (cAMP) induces apical chloride (Cl^-^) secretion from epithelial cells through cystic fibrosis transmembrane conductance regulator (CFTR).^1^ The resulting ionic gradient draws Na^+^ and then water out of the tissue and into the lumen, where it hydrates mucus and can contribute to epithelial “flushing”.^2, 3^ Loss-of-function mutations in CFTR cause cystic fibrosis, a disease characterized by viscous mucus, reduced lung function, and bacterial overgrowth.^1^ Epithelial Cl^-^/water secretion can also occur via calcium-dependent ion channels, with muscarinic acetylcholine receptors (mAChR) in the basolateral membrane often inducing the necessary intracellular calcium flux.^2^ Acetylcholine (ACh) is a canonical neurotransmitter synthesized by the enzyme choline acetyltransferase (*Chat*). Neurons innervating mucosal barriers can induce ACh-dependent fluid secretion, but non-neuronal sources of ACh have now been widely reported in other contexts.^3–7^

Among epithelial cells, tuft cells are the dominant source of ACh.^8, 9^ Found across mucosal tissues, they are a lineage of chemosensory cells that monitor the lumenal microenvironment and release effectors to regulate the mucosa. *Chat* expression is part of a transcriptional signature shared by all murine tuft cells^10, 11^ and ChAT protein has been detected in human tuft cells in the intestine and airways.^12, 13^ The function of tuft cell-derived ACh has also been studied in several tissues. For example, tuft cells in the nasal epithelium sense bitter and bacteria-derived ligands^14^ and secrete ACh, which signals on neurons to induce neurogenic inflammation.^15^ Tracheal tuft cells activate nicotinic ACh receptors (nAChRs) to cause a brief cessation in breathing^16^ and mAChRs on neighboring epithelial cells to increase ciliary beat frequency^8, 17^ in response to similar ligands. Likewise, tuft cells in the urethra use ACh to activate neurons and regulate urine release.^18, 19^ However, the function of tuft cell-derived ACh in the intestine is unknown, nor has a link between tuft cells and fluid secretion been tested.

Small intestinal (SI) tuft cells play a critical role in the initiation of “Type 2” immune responses to helminth infection and colonization by *Tritrichomonas sp.* protists.^20–22^ Tuft cells express SUCNR1, the receptor for extracellular succinate, which *Tritrichomonas sp.* and the microbiota secrete as a metabolite.^11, 23, 24^ SUCNR1 signaling causes intracellular Ca^2+^ flux that opens the cation channel TRPM5. The resulting Na^+^ influx depolarizes the tuft cell and likely regulates secretion of most tuft cell effector molecules.^25, 26^ *Sucnr1^-/-^* mice fail to detect *Tritrichomonas sp.* colonization, but the immune response to helminth infection is unaffected.^11, 24^ Nonetheless, sensing of both helminths and protists is severely attenuated in *Trpm5^-/-^* mice or *Pou2f3^-/-^* mice that lack tuft cells entirely.^11, 20, 22^

Once activated by lumenal signals, tuft cells produce IL-25 and, in some cases, leukotriene C4 (LTC_4_) to activate resident group 2 innate lymphoid cells (ILC2s) in the underlying *lamina propria* (LP).^21, 27^ ILC2s secrete canonical Type 2 cytokines, including IL-13, that collectively recruit Type 2 immune cells and coordinate intestinal remodeling. Among its many targets, IL-13 produced by ILC2s signals on undifferentiated epithelial cells to bias differentiation towards mucus-producing goblet cells and tuft cells.^20, 21, 28, 29^ Given the 3-5 day turnover of the intestinal epithelium, this feed-forward process, known as the tuft-ILC2 circuit, results in dramatic hyperplasia of both goblet cells and tuft cells, the latter of which increase 10-fold.^20–22^

The Type 2 effector functions that clear worms from the intestine,^30, 31^ collectively referred to as “weep and sweep,” require the coordination of multiple signals. In addition to increasing the number of tuft and goblet cells, IL-13 upregulates production of mucus and anti-helminthic/microbial peptides in the epithelium,^32–34^ increases fluid secretion,^35^ and increases expression of mAChRs on smooth muscle,^36–38^ but actual secretion (“weep”) and muscle contraction (“sweep”) generally require additional signals. ACh is one molecule that can acutely activate both weep and sweep,^38^ and mAChRs are required for helminth clearance.^39, 40^ Conversely, helminths secrete ACh esterases (AChE), likely in an attempt to inhibit weep and sweep responses.^41–43^ The sources of ACh in Type 2 immunity, however, are not defined.

By sensing lumenal signals and activating ILC2s, tuft cells serve as sentinels for intestinal Type 2 immunity, but the fact that many more tuft cells are generated after the agonist has been detected suggests an additional effector function for these cells. Here we describe such an effector function, which is independent of ILC2s. We show that in response to sensing of succinate or direct activation of TRPM5, tuft cells secrete ACh to induce epithelial fluid secretion in the intestine and airways. During Type 2 tissue remodeling, *Chat*+ tuft cells increase in number, enhancing the fluid secretion response. Upon helminth infection, mice with *Chat*-deficient tuft cells experience delayed helminth clearance despite normal tuft-ILC2 circuit activation. We conclude that tuft cell-derived ACh regulates epithelial fluid secretion, and that this effector function can contribute to Type 2 immune responses during helminth infection.

## Results

### SI tuft cells express Chat in a proximal to distal gradient

Neuronal *Chat* is important for intestinal function, but the role of *Chat* in intestinal tuft cells has not been studied. To assess *Chat* expression by tuft cells in the SI at single cell resolution, we employed *Chat-GFP* transgenic reporter mice. Immunolabeling for GFP colocalized with the tuft cell marker DCLK1 in both the proximal SI (pSI; first 5-10 cm) and distal SI (dSI; last 5-10 cm) (Fig. 1A). By flow cytometry, >99% of GFP*+* epithelial cells stained for the tuft cell-specific *Il25-RFP* reporter (Fig. 1B). However, not all tuft cells were GFP+ (Fig. S1A), and we observed a gradient in the frequency of GFP+ tuft cells that increased from 40% of all tuft cells in the pSI to 80% in the dSI (Fig. 1C-D). The discovery of GFP*-*negative tuft cells was unexpected, as the *Chat* reporter marks nearly 100% of tuft cells in other tissues.^8^ To validate our findings, we crossed *Chat-*Cre mice, in which Cre is expressed from the endogenous *Chat* locus, to *Rosa26::STOP^fl/fl^::CAG-tdTomato* (Ai9) mice for lineage tracing. Using CD24 and Siglec-F to identify tuft cells, we again observed both reporter-positive and -negative tuft cells, with an increased frequency of reporter-positive tuft cells in the distal compared to the proximal SI (Fig. S1B).

**Figure 1:**
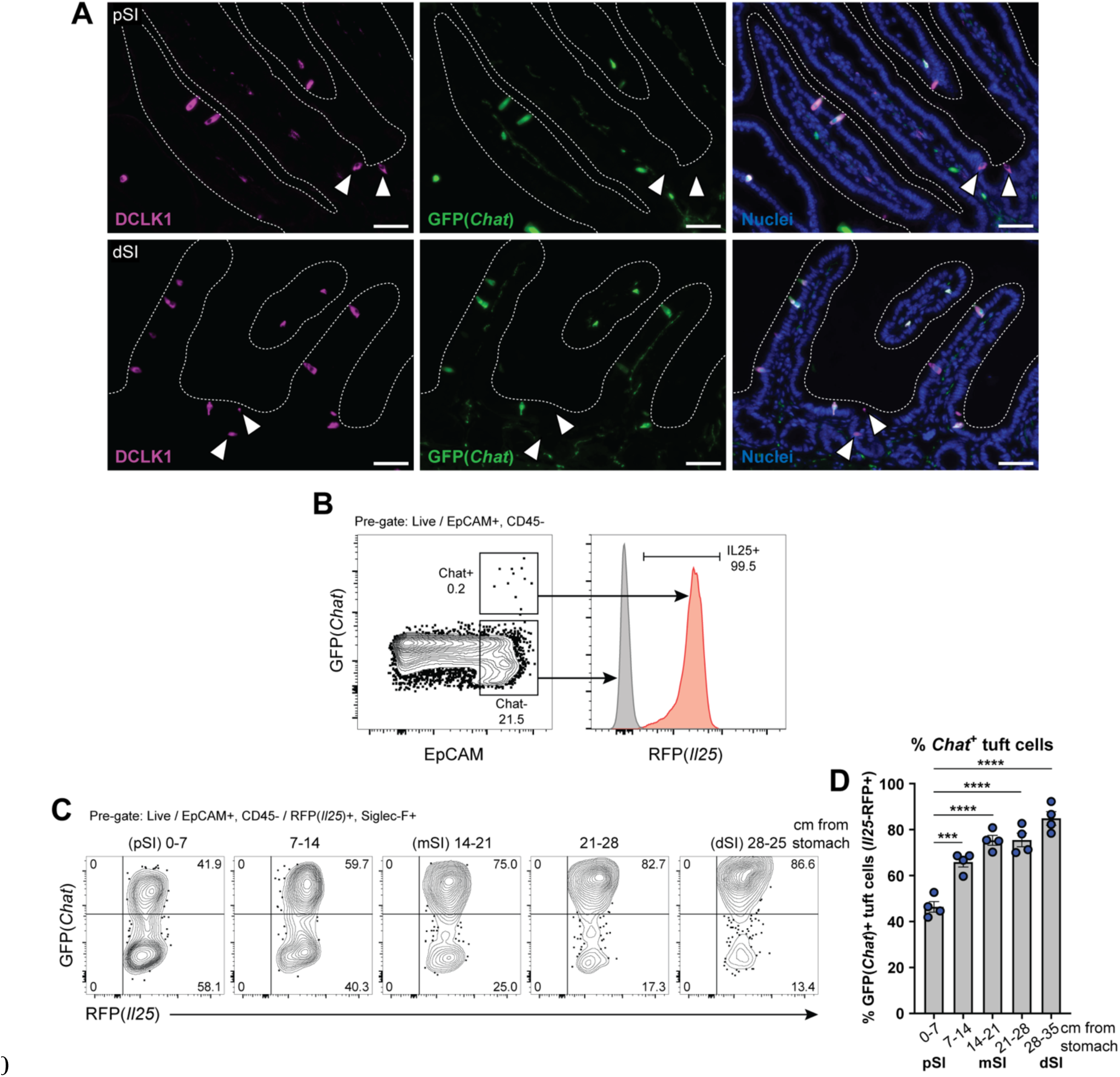
SI tuft cells express *Chat* in a proximal to distal gradient. (**A**) Representative images of GFP(*Chat*) expression (green) by DCLK1+ tuft cells (magenta) in the proximal SI (pSI) and distal SI (dSI) by immunofluorescence. White arrows indicate GFP(*Chat*)-tuft cells. Nuclei stained with DAPI (blue). Scale bars: 50 µm (**B**) GFP+ epithelial cells (EpCAM+) are RFP(*Il25*)+ tuft cells. (**C** and **D**) (C) Representative flow cytometry and (D) quantification of the percentage of GFP+ tuft cells by sequential 7 cm section across the length of the SI. In D, each symbol represents an individual mouse (columns represent different tissues from same mouse) from three pooled experiments. *p < 0.05, **p < 0.01, ***p < 0.001 by one way ANOVA with Tukey’s multiple comparisons test (D). mSI, medial SI. Graphs depict mean +/-SEM. Also see Figure S1.

Given the binary nature of *Chat-GFP* expression in SI tuft cells, we hypothesized that *Chat* might mark transcriptionally distinct tuft cell subsets. We therefore sorted GFP*+* and GFP-tuft cells, performed bulk RNA sequencing, and identified differentially expressed genes (DEGs) (Fig. S1C, Table S1, Data File S1). Surprisingly, despite the binary nature of GFP expression, *Chat* was downregulated only 2.8-fold (FDR = .0009) in GFP-cells, suggesting translation is regulated via untranslated regions of the endogenous *Chat* transcript that are retained in the transgene (Fig. S1C). More broadly, even with a lenient fold-change (FC) cutoff of 2 (FDR <.01), there were only 105 DEGs. None of the downregulated genes were part of the SI tuft cell signature,^44^ but *Sucnr1* (FC = 3.3, FDR = .0007) and the downstream G alpha subunit *Gnat3* (FC = 4.9, FDR = .0001) were upregulated in GFP+ cells (Fig S1C, Table S1). Comparing *Chat+* tuft cells to previously reported intestinal tuft cell subsets “Tuft-1” and “Tuft-2”,^9^ we found greater enrichment for Tuft-2 genes (Fig. S1D), but the best match was with a dSI tuft cell signature we generated by sorting tuft cells from the proximal and distal 5 cm of unmanipulated intestines (Fig. S1D-E, Table S2, Data File S2). This signature similarly includes *Sucnr1* and *Gnat3,* and we hypothesize that transcriptional differences between GFP*+* and GFP*-* cells resulted mostly from a distal bias among sorted GFP*+* cells, consistent with the gradient we observed (Fig. 1D).

We also considered the possibility that *Chat*-cells were an immature stage before *Chat+* cells, but while GFP+ cells predominated in the villi and GFP-cells in the crypts, there were still GFP*+* cells in the crypts (Fig. S1F) and GFP-cells at the villus tips (Fig. S1G), making a developmental relationship unlikely. Both IL4ra-dependent and IL4ra-independent tuft cells have been identified in the SI,^45^ but we found many *Chat-GFP*+ tuft cells in the SI of *Il4ra^-/-^* mice, suggesting *Chat* expression is not exclusive to IL-13-induced tuft cells (Fig. S1H). The mechanisms regulating *Chat* expression in tuft cells therefore remain unknown.

**Supplemental Figure 1:**
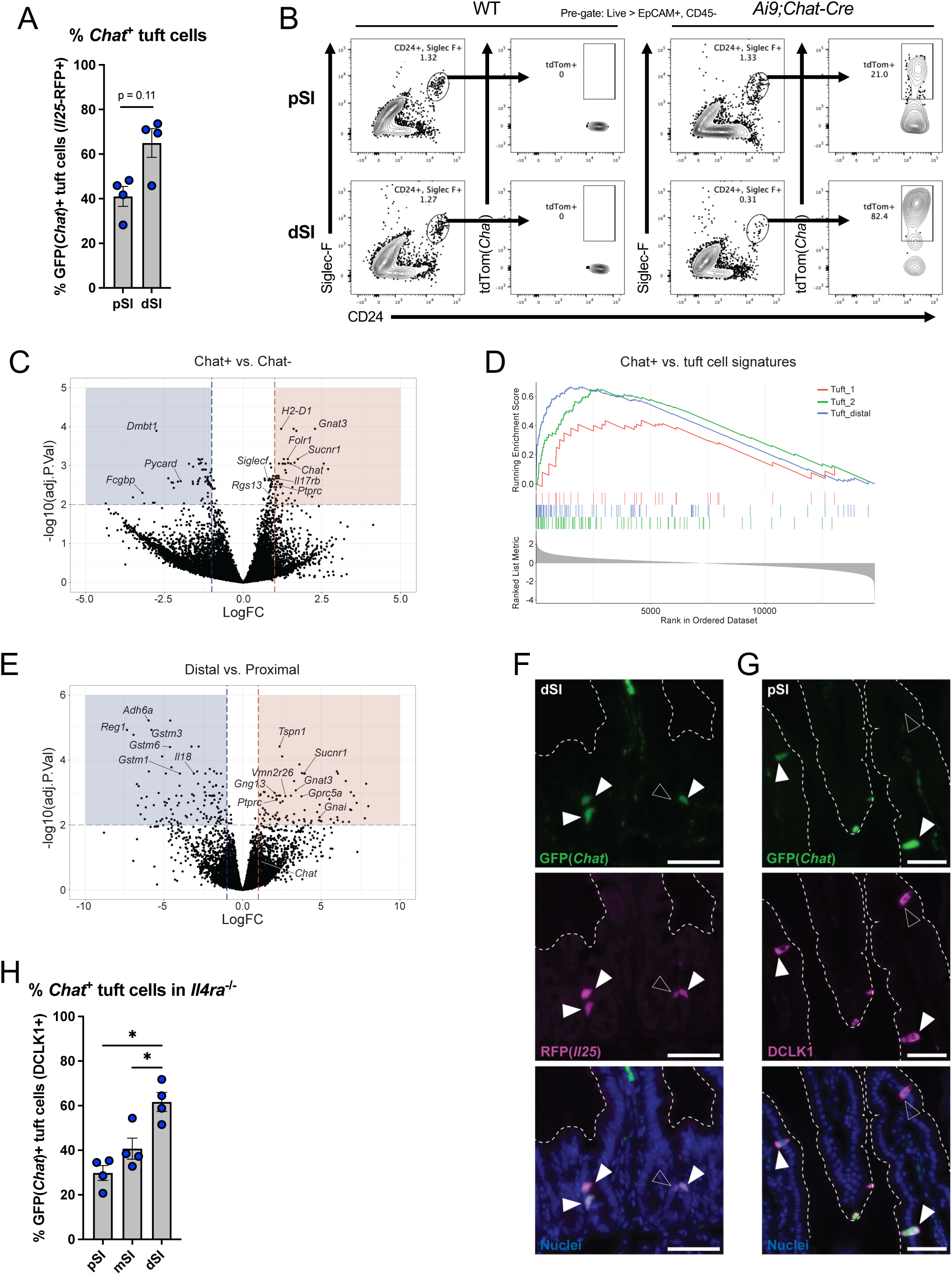
(**A**) Quantification of GFP+ tuft cells (RFP+) from pSI and dSI of *Chat-GFP*;*Il25*^RFP/+^ mice by immunofluorescence (IF). (**B**) Representative flow cytometry of traced tdTom+ tuft cells (CD24+, Siglec-F+) from pSI and dSI of wild-type (WT) and *Ai9;Chat-Cre* mice. (**C**) Volcano plot showing log2FC of genes expressed in *Chat*+ tuft cells (n=4) versus *Chat-* tuft cells (n=3) sorted from whole SI of *Chat-GFP*;*Il25*^RFP/+^ mice. (**D**) Gene set enrichment analysis comparing *Chat+* tuft cell gene expression to Tuft-1 and Tuft-2 consensus gene signatures and the dSI tuft cell signature from (**E**) Volcano plot showing log2FC of genes expressed in tuft cells sorted from the dSI (n=4) versus tuft cells sorted from the pSI (n=4) of B6 mice. (**F**) Representative immunofluorescence image showing GFP+ (green) tuft cells (RFP+, magenta) in the SI crypt (solid white arrows), next to one GFP-tuft cell (open white arrow). Nuclei stained with DAPI (blue). Scale bars: 50 µm. (**G**) Representative immunofluorescence image showing a GFP-(green) tuft cell (DCLK1+, magenta) at the villus tip (open white arrow), past other GFP+ tuft cells (solid white arrows). Nuclei stained with DAPI (blue). Scale bars: 50 µm. (**H**) Quantification of GFP(*Chat*)+ tuft cells (DCLK1+) from denoted tissues of *Il4ra*^-/-^*;Chat-GFP* mice by immunofluorescence. In the graphs, each symbol represents an individual mouse (columns represent different tissues from same mouse) from two pooled experiments. *p < 0.05, **p < 0.01, ***p < 0.001 by Mann-Whitney test (A) or one way ANOVA with Tukey’s multiple comparisons test (G). mSI, medial SI. Graphs depict mean +/-SEM.

### ACh from tuft cells induces fluid secretion from the SI epithelium

ACh rapidly induces fluid, mucus, and antimicrobial peptide secretion when it binds muscarinic ACh receptors (mAChRs) on SI epithelial cells.^4, 46–48^ Classically, enteric neurons are considered the primary source of ACh that regulates SI secretion,^4, 49, 50^ but we hypothesized that tuft cells could link lumenal chemosensing to epithelial secretion via basolateral release of ACh.

To make sensitive, real-time measurements of SI epithelial electrophysiology, we employed Ussing chambers,^51^ which have been used to measure ACh-induced epithelial ion flux.^52^ Two chambers containing physiologic buffer are separated by a piece of SI epithelium and the voltage across the epithelium is clamped. When negatively charged ions (e.g. Cl^-^) are secreted into the lumenal chamber, current is injected into the basolateral chamber to restore the voltage (Fig. 2A). This “short-circuit” current, or Isc, is directly proportional to ion flux.^51^

**Figure 2:**
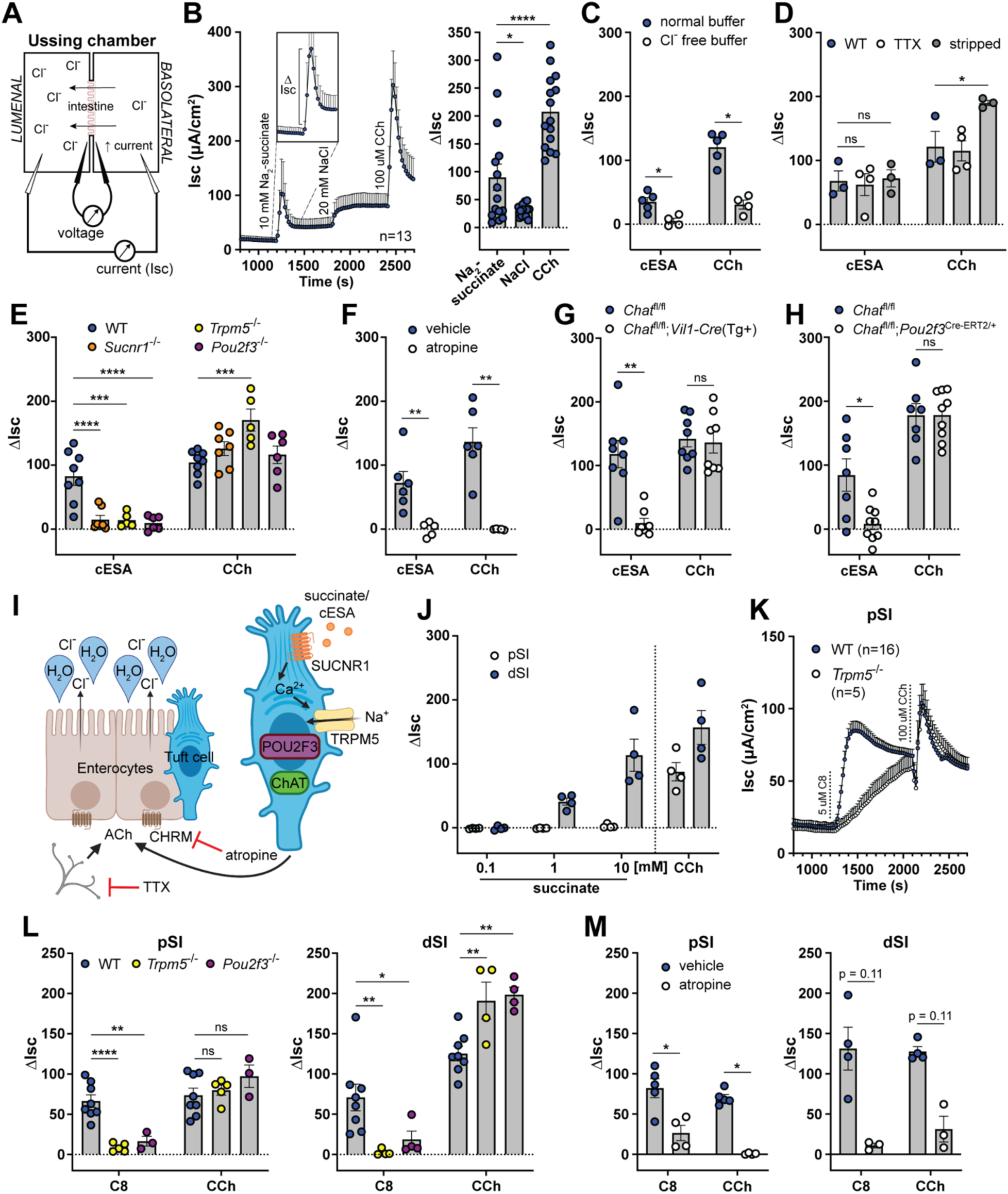
Tuft cell-derived ACh induces epithelial fluid secretion. (**A**) Ussing chamber schematic. (**B**) Average Isc traces and quantification of the delta Isc (Λ1Isc, see inset and bar graph) of WT dSI tissue stimulated as indicated (10 mM Na2-succinate and 20 mM NaCl, lumenal; 100 µM CCh, basolateral). (**C**) ΔIsc values of WT dSI in presence of normal chloride-(Cl^-^) containing buffer or buffer selectively lacking Cl^-^, stimulated as indicated (10 mM cESA, lumenal). (**D**) ΔIsc values of WT intact dSI compared to stripped dSI and dSI pretreated 15 min with TTX (1 µM, basolateral), stimulated as indicated. (**E**) ΔIsc values of dSI from mice of indicated genotypes stimulated as indicated. (**F**) ΔIsc values of WT dSI compared to dSI pretreated 15 min with pan-CHRM inhibitor atropine (10 µM, basolateral), stimulated as indicated. (**G** and **H**) ΔIsc values of dSI with (G) epithelial cell-(*Vil1-Cre*) and (H) tuft cell-specific (*Pou2f3^ERT^*^2^*^-Cre/+^*) *Chat* deletion, stimulated as indicated. (**I**) Model of tuft cell chemosensing of succinate driving ACh-dependent fluid secretion independent of neurons. (**J**) ΔIsc values of WT pSI and dSI stimulated as indicated. (**K**) Average Isc traces of pSI from WT or *Trpm5*^-/-^ mice stimulated as indicated (5 µM C8, basolateral). (**L** and **M**) ΔIsc values of WT tissues compared to (L) tissues from indicated genotypes or (M) tissues pretreated 15 min with atropine (10 µM, basolateral), stimulated as indicated. In the graphs, each symbol represents an individual mouse (one tissue or average of two) pooled from two or more experiments. Groups represent sequential stimulations of the same tissue. *p < 0.05, **p < 0.01, ***p < 0.001, ****p < 0.001 by RM one way ANOVA with Tukey’s multiple comparisons test (B), two way ANOVA with Dunnett’s multiple comparisons test (D, E, L), multiple Mann-Whitney tests with Holm Sídák’s multiple comparisons test (C, F, G, H, M). ns, not significant. Graphs depict mean +/-SEM. Also see Figure S2.

To test whether tuft cells regulate ion flux, we mounted SI tissue from unmanipulated mice in the Ussing chamber and stimulated with Na_2_-succinate (succinate), since it is the best-defined ligand for SI tuft cells.^11, 23, 24^ When we stimulated the lumenal side of the SI tissue with 10 mM succinate, we recorded a rapid increase in the Isc that lasted several minutes before returning close to baseline (Fig. 2B). We quantified this response by measuring the change in Isc from the baseline before stimulation to its peak, known as the delta Isc (ι1Isc; see Fig. 2B inset). To control for the addition of sodium, we tested NaCl at an equimolar concentration of sodium (20 mM). NaCl was sufficient to increase the Isc, but the response was significantly smaller than the succinate response and failed to return to baseline. The succinate response, on the other hand, had similar kinetics to the response elicited by the ACh mimic carbachol (CCh), given basolaterally to maximally stimulate mAChRs (Fig. 2B). The succinate response was greatly diminished in SI from *Sucnr1^-/-^* mice, though a residual, likely sodium-dependent increase remained (Fig. S2A). To avoid the sodium response, we used the synthetic SUCNR1 agonist cis-epoxysuccinic acid (cESA),^53^ which stimulated an Isc response comparable to succinate, but that was entirely *Sucnr1-*dependent (Fig. S2A). We used cESA in place of succinate for most of the subsequent Ussing experiments.

To further characterize the succinate/cESA Isc response, we began by testing the role of epithelial polarity. cESA induced ion flux when given lumenally, but not basolaterally, consistent with lumenal restriction of SUCNR1 in tuft cells (Fig. S2B). Conversely, CCh stimulated ion flux when given basolaterally, but not lumenally, consistent with basolateral restriction of mAChRs on intestinal epithelial cells (IECs).^4^ Since the Isc represents net ion flux across the epithelium, the increased Isc response to succinate could be due to either increased lumenal secretion of negatively charged anions (e.g. Cl^-^, HCO3^-^) or increased absorption of positively charged cations (e.g. Na^+^, K^+^) from the lumen.^4^ To test the contribution of Cl^-^ ions selectively, we replaced Cl^-^ with gluconate, which cannot cross the epithelium,^51, 54^ and found that both the cESA and CCh responses were abrogated (Fig. 2C). Bumetanide, an inhibitor of the basolateral chloride transporter NKCC1 that is required for sustained Cl^-^ secretion,^1^ likewise decreased the response to cESA (Fig. S2C).

Since enteric neurons are both a major source of ACh and major regulators of fluid secretion in the intestine, we investigated the possibility that SI tuft cells activate enteric neurons. Studies of *Chat*+ tuft cells in the airways have emphasized their close contact with neurons and provided evidence that signaling can occur from tuft cells to neurons, often via release of ACh.^15, 16^ We therefore looked for similar tuft-neuronal connections by microscopy using the *Chat*-*GFP* reporter, which marks cholinergic intestinal neurons as well as tuft cells. We found some instances where a GFP+ tuft cell was approached by a GFP+ neuron, but we never saw neurons extend into the epithelium and contact tuft cells, as they do in the airways (Fig. S2D). *Chat*+ neurons represent only a subset of total intestinal neurons,^55^ yet staining for the pan-neuronal marker BIII tubulin (TUJ1) revealed no additional neuronal contacts (Fig. S2E), suggesting that neuronal contacts with tuft cells, much less “synapses”, are uncommon in the SI.

Recognizing that signaling can occur without direct contact, we experimentally tested the requirement for neurons in the succinate response. First, we disrupted neuronal integrity by physically “stripping” the submucosa off the back of the epithelium to eliminate most of the submucosal and all of the myenteric neuronal plexuses.^51^ Alternatively, we used tetrodotoxin (TTX) to inhibit neuronal action potentials in intact SI tissue.^4^ Neither treatment reduced the cESA or CCh responses; in the stripped tissue the CCh response was instead increased, likely due to enhanced diffusion (Fig. 2D). Altogether, our data show that succinate/cESA binds to SUCNR1 expressed apically on epithelial cells and induces chloride-dependent fluid secretion, independently of enteric neurons or submucosal tissue.

To confirm that tuft cells were the cells that sensed succinate/cESA to initiate the secretion response, we stimulated dSI from tuft cell-deficient *Pou2f3^-/-^*, and chemosensing-deficient *Trpm5^-/-^*mice. As with SI tissue from *Sucnr1^-/-^*mice, tissues from these mice failed to respond to cESA (Fig. 2E). Importantly, the CCh response was intact in all knockout mice, demonstrating that the tissue’s capacity for ACh-dependent fluid secretion was unaltered. To test whether ACh was involved in the cESA response, we pretreated the dSI with the pan-mAChR inhibitor atropine and found that this completely blocked the cESA and CCh responses (Fig. 2F). Deletion of *Chat* from IECs using *Chat^fl/fl^;Vil1-Cre(Tg+)* mice also abrogated the cESA response (Fig. 2G). Although tuft cells are the only *Chat*-expressing IECs, we also deleted *Chat* in tuft cells specifically using *Chat^fl/fl^;Pou2f3^Cre-ERT^*^2^*^/+^* mice, and confirmed that tuft cell-derived ACh production was required for cESA-induced fluid secretion (Fig. 2H).

Other tuft cell effectors (e.g. LTC_4_ or PGD_2_) have been implicated in acute responses and tuft cells themselves express the receptor for IL-25,^11, 56, 57^ but we excluded the involvement of IL-25 and LTC_4_ in ion flux using dSI from *Il25^-/-^* and *Alox5^fl/fl^;Vil1-Cre1000(Tg+)* mice, respectively (Fig. S2F,G). Pretreatment with the COX inhibitor ibuprofen to block PGD_2_ synthesis also did not affect the cESA response (Fig. S2H). Furthermore, PGD_2_, which has not been previously linked with fluid secretion but has been reported to induce mucus secretion from goblet cells (GCs) in the colon,^56^ did not induce ion flux in the dSI (or the colon) when administered basolaterally (Fig. S2I). We also investigated the possibility that tuft cells were signaling to neighboring cells via gap junctions.^58^ The gap junction inhibitor carbenoxolone partially blocked cAMP-driven fluid secretion induced by IBMX + forskolin, but had no effect on cESA or CCh responses (Fig. S2J). Thus, we have demonstrated that tuft cells in the dSI sense lumenal succinate/cESA and release ACh basolaterally, which stimulates mAChR-dependent Cl^-^ ion secretion (Fig. 2I). The response is epithelium-intrinsic and does not involve enteric neurons.

**Supplemental Figure 2:**
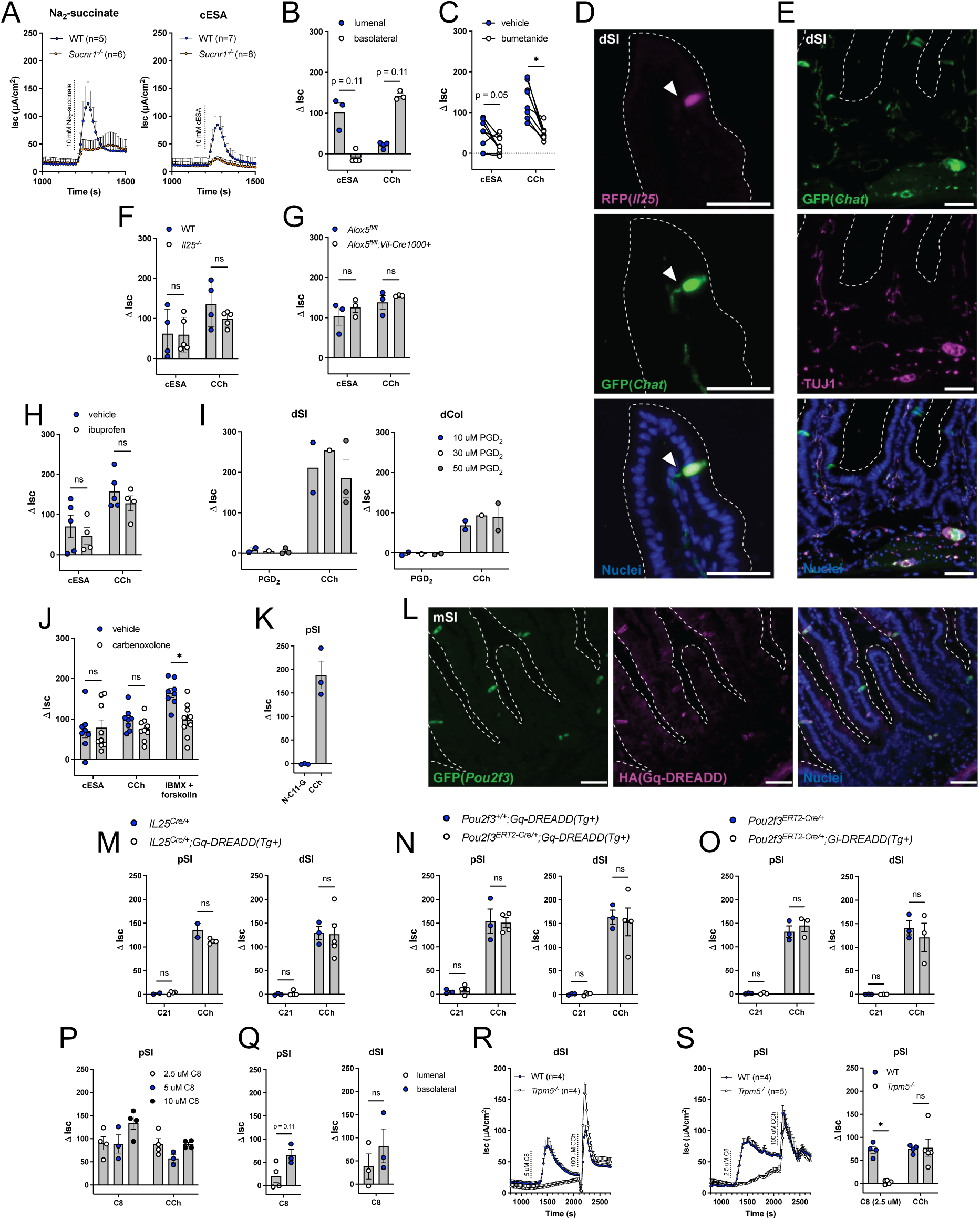
(**A**) Average Isc traces of dSI from WT or *Sucnr1*^-/-^ mice stimulated as indicate (Na2-succinate and cESA, lumenal). (**B**) ΔIsc values of WT dSI stimulated as indicated. (**C**) ΔIsc values of WT dSI pretreated 15 min with vehicle or bumetanide (100 µM, basolateral), stimulated as indicated (10 mM cESA lumenal; 100 µM CCh, basolateral). (**D**) Representative immunofluorescence image of a GFP*+* (green) neuronal process (indicated by solid white arrow) approaching a GFP+/RFP+ (magenta) tuft cell from the dSI of *Chat-GFP*;*Il25^RFP/+^* mice. Nuclei stained with DAPI (blue). Scale bars: 50 µm. (**E**) Representative immunofluorescence image of GFP*+* (green) neurons co-stained for BIII tubulin (TUJ1, magenta) in the dSI. Nuclei stained with DAPI (blue). Scale bars: 50 µm. (**F**, **G**, and **H**) (F and G) ΔIsc values of dSI from indicated genotypes or (H) WT dSI compared to dSI pretreated 15 min with ibuprofen (10 µM, bilateral), stimulated as indicated. (**I**) ΔIsc values of WT tissues stimulated as indicated (PGD2, basolateral). (**J**) ΔIsc values for WT dSI compared to dSI pretreated 15 min with carbenoxolone (1 mM, lumenal), stimulated as indicated (100 µM IBMX + 10 µM forskolin, bilateral). (**K**) ΔIsc values of WT pSI stimulated as indicated (100 µM N-C11-G, lumenal). (**L**) Representative immunofluorescence image of GFP+ (green) tuft cells expressing HA+ Gq-DREADDs (magenta) in the crypts and villi of the medial SI (mSI). Nuclei stained with DAPI (blue). Scale bars: 50 µm. (**M**, **N**, and **O**) (M) ΔIsc values of indicated tissues from unmanipulated mice or (N and O) indicated tissues from mice 7 days after start of tamoxifen chow, stimulated as indicated (1 µM C21, bilateral). (**P**) ΔIsc values of WT pSI stimulated as indicated (C8, bilateral). (**Q**) ΔIsc values of WT tissues from pSI and dSI stimulated as indicated. (**R**) Average Isc traces of dSI stimulated as indicated (5 µM C8, basolateral). (**S**) Average Isc traces and ΔIsc values of pSI stimulated as indicated (2.5 µM C8, basolateral). In the graphs, each symbol represents an individual mouse (one tissue or average of two) pooled from two or more experiments. Groups represent sequential stimulations of the same tissue. In (C) paired vehicle and bumetanide-treated tissues are from the same mouse. *p < 0.05, **p < 0.01, ***p < 0.001, ****p < 0.001 by multiple Mann-Whitney tests with Holm Sídák’s multiple comparisons test (B, F-H, J, M-O, S), Wilcoxon matched-pairs signed rank test with Holm Sídák’s multiple comparisons test (C), or Mann-Whitney test (Q). ns, not significant. Graphs depict mean +/-SEM.

### A TRPM5 agonist induces tuft- and ACh-dependent fluid secretion in the pSI and dSI

In characterizing the succinate response, we found that the pSI did not respond to succinate stimulation (Fig. 2J). This finding is consistent with greater succinate receptor (*Sucnr1*) expression in tuft cells from the dSI compared to the pSI (Table S2),^23^ and may also reflect the reduced frequency of *Chat*+ tuft cells in the pSI (Fig. 1D). Since the pSI was responsive to CCh, we reasoned that pSI tuft cells could induce fluid secretion if properly stimulated. We first tested several putative tuft cell ligands. Worm excretory and secretory products from *Nippostrongylus brasiliensis* (*Nb*), known as NES, failed to stimulate ion flux whether made from infective L3 larvae or adult worms (data not shown). The bacterial metabolite N-C11-G did not induce ion flux either (Fig. S2K). Next, we tried a chemogenetic approach with Gq-or Gi-coupled receptors that respond only to synthetic ligands (DREADDs)^59^ using constitutive (*Il25-Cre*) or inducible (*Pou2f3^Cre-ERT^*^2^*^/+^*) for tuft cell-specific expression. Although tuft cells expressed the HA tag included in the DREADD constructs, stimulation with Compound 21^60^ was insufficient to drive a fluid secretion response in the pSI or dSI (Fig. S2L-O). Perhaps G proteins required for DREADD function are not available in tuft cells or the signals induced downstream of DREADD activation are not sufficient to induce ACh release.

Finally, we decided to stimulate TRPM5 directly, since all tuft cell chemosensing pathways identified to date converge on TRPM5. We acquired a TRPM5 agonist compound called Class 8 (C8),^61, 62^ and found that it induced ion flux in both the pSI and dSI when administered lumenally or basolaterally, with a trend toward higher basolateral responses (Fig. 2K, S2P-Q). The C8 response was similar to cESA- or CCh-induced ion flux, and was TRPM5-dependent in the pSI and dSI (Fig. 2K-L, S2R). The C8 response was also tuft cell- and ACh-dependent in both locations (Fig. 2L-M). In the pSI, C8 induced a slow TRPM5-independent increase in Isc (Fig. 2K), but this off-target effect could be eliminated by lowering the dose of C8 (Fig. S2S). We conclude that in response to direct TRPM5 activation, tuft cells in the pSI and dSI can release ACh to induce fluid secretion from the intestinal epithelium.

### Tuft cell-mediated fluid secretion occurs across mucosal tissues and is detectable in vivo

We previously found that *Sucnr1* expression is even higher in tracheal tuft cells than those of the SI,^11^ so to test if tuft cells regulate fluid secretion at multiple mucosal barriers, we stimulated tracheal tissue in the Ussing chamber with succinate. As in the dSI, we found that this induced a rapid increase in the Isc that was *Pou2f3-*, *Sucnr1-*, and *Trpm5*-dependent (Fig. 3A-B). In addition, the response was mAChR-dependent (Fig. 3C). By comparison, the cecum and colon, where tuft cells express *Sucnr1* at low levels,^11^ responded only weakly to succinate stimulation and in a tuft-independent manner (Fig. S3A). Colonic tissue did respond to TRPM5 agonism with C8, with a larger response in the proximal colon (pCol) than the distal colon (dCol) (Fig. 3D-E). The reported tuft ligand N-C11-G^56^ did not stimulate fluid secretion from the dSI or dCol (Fig. S3B), and also failed to elicit tuft-dependent leukotriene production from intestinal epithelial monolayers (Fig. S3C). Therefore, tuft cell control of epithelial fluid secretion is a common effector function across barrier tissues.

**Figure 3:**
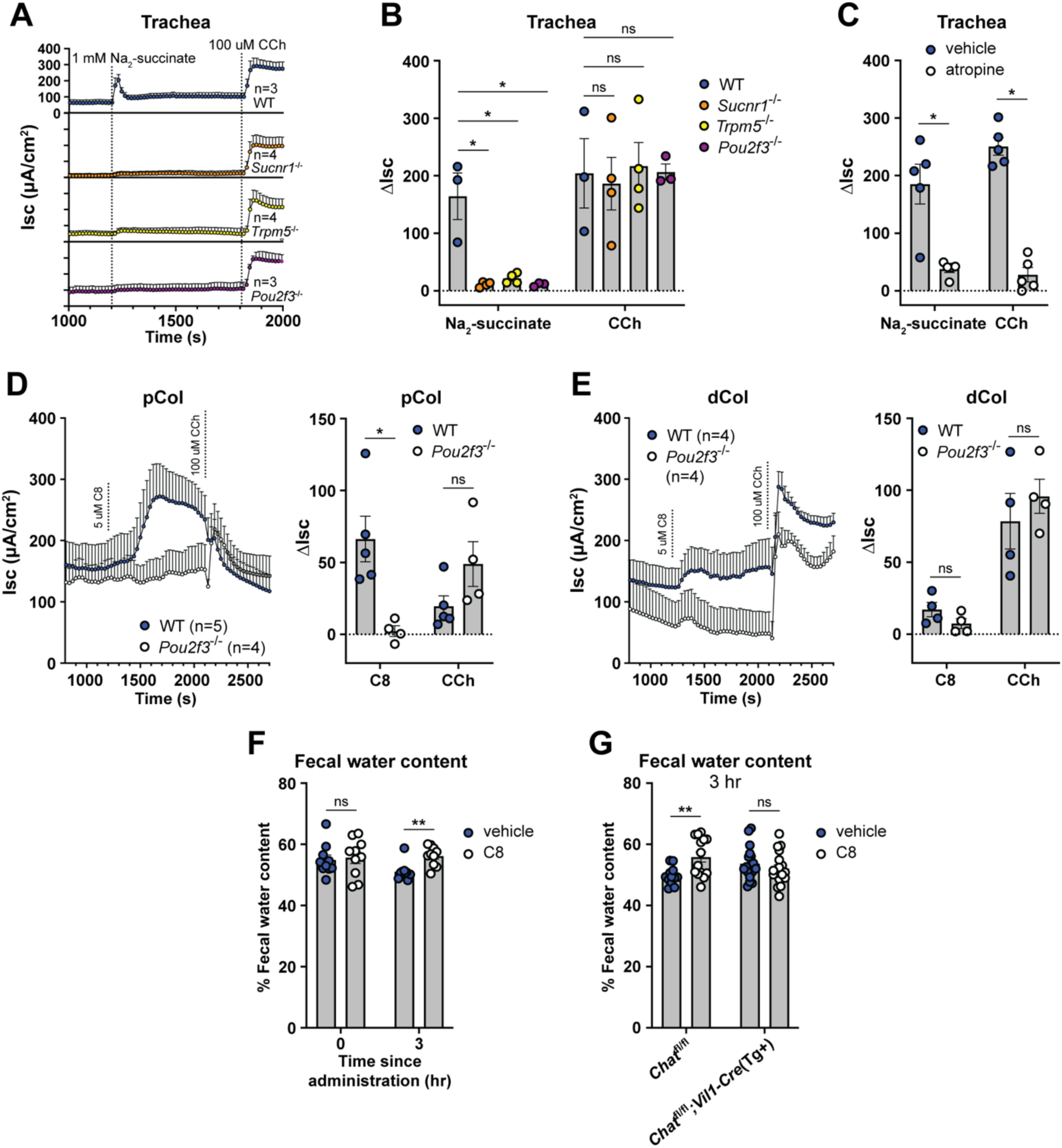
Tuft cell-mediated fluid secretion occurs across mucosal tissues and is detectable in vivo. (**A** and **B**) (A) Average Isc traces and (B) ΔIsc values of trachea from mice of indicated genotypes stimulated as indicated (1 mM Na2-succinate, lumenal; 100 µM CCh, basolateral). (**C**) ΔIsc values of WT trachea compared to trachea pretreated 15 min with atropine (10 µM, basolateral), stimulated as indicated. (**D** and **E**) Average Isc traces and ΔIsc values of WT and *Pou2f3*^-/-^ tissues stimulated as indicated (5 µM C8, basolateral). (**F** and **G**) Quantification of water content of fecal pellets collected from (A) WT or (B) *Chat-fl;Vil1-Cre(Tg+)* mice treated orally with vehicle or C8 (30 mg/kg) for the indicated durations. In the graphs, each symbol represents an individual mouse (one tissue or average of two) pooled from two or more experiments. Groups represent sequential stimulations or timepoints of the same tissue or animal. *p < 0.05, **p < 0.01, ***p < 0.001, ****p < 0.001 by, two way ANOVA with Tukey’s multiple comparisons test (B), or multiple Mann-Whitney tests with Holm Sídák’s multiple comparisons test (C-G). ns, not significant. Graphs depict mean +/-SEM. Also see Figure S3.

The Ussing chamber measures ion flux but cannot measure water movement directly. We therefore wanted to test if activating tuft cells *in vivo* could induce fluid secretion into the intestine. We dosed mice with C8 or vehicle in the morning and then measured the wet weight of fecal pellets 3 hours later. In mice given vehicle, the fecal water content declined, likely due to reduced water intake during the day. C8 treatment prevented this decline, indicating sustained fluid secretion (Fig. 3F). Importantly, this fluid secretion was dependent on epithelial *Chat* (Fig. 3G, S3D). Thus, activation of tuft cells along the intestinal tract induces ACh-dependent ion flux that drives fluid secretion into the intestinal lumen.

**Supplemental Figure 3:**
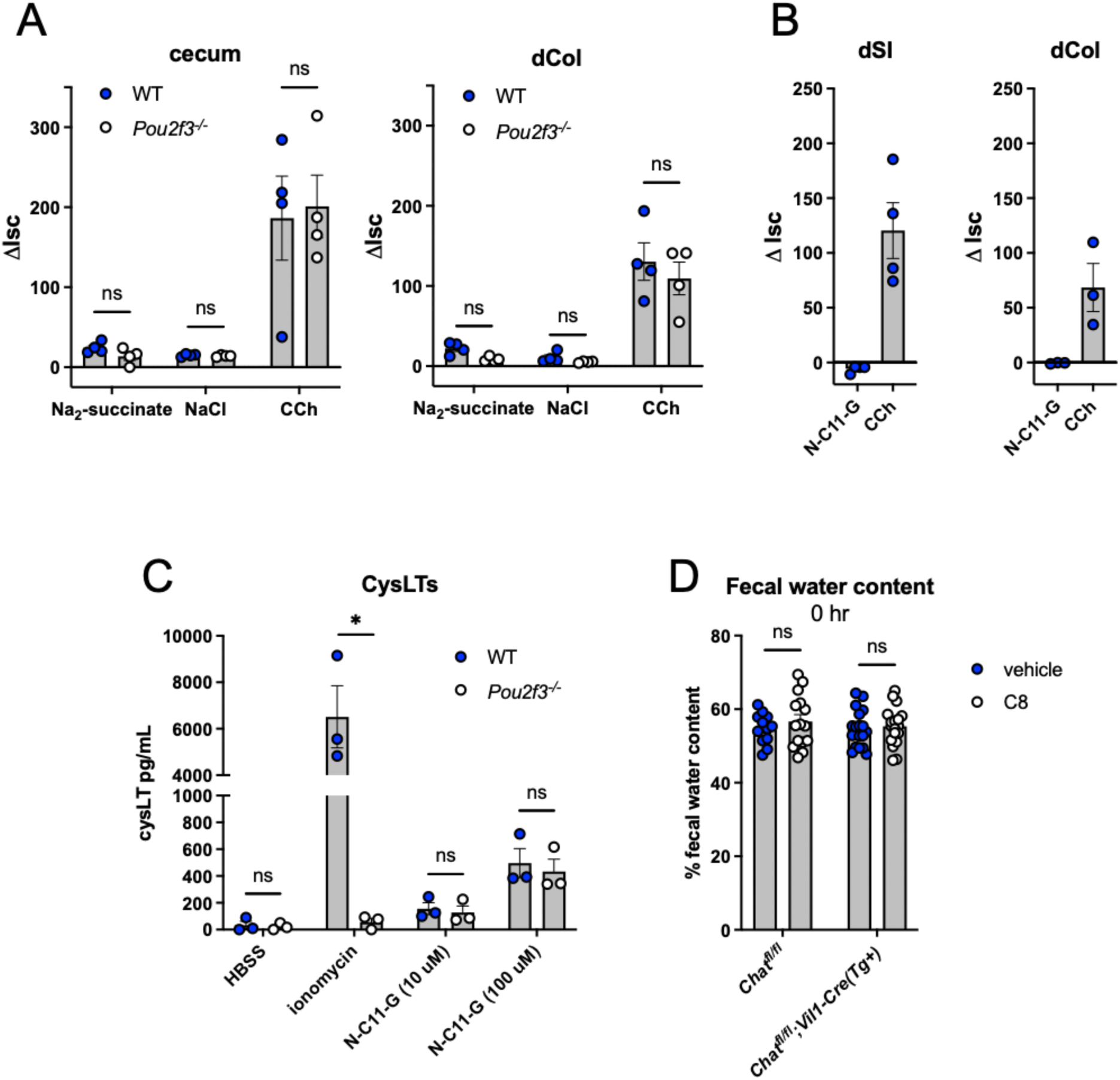
(**A**) ΔIsc values of WT and *Pou2f3*^-/-^ tissues stimulated as indicated (10 mM Na2-succinate and 20 mM NaCl, lumenal; 100 µM CCh, basolateral). (**B**) ΔIsc values of WT tissues stimulated as indicated (100 µM N-C11-G, lumenal). (**C**) Cysteinyl leukotriene (CysLTs) production from WT and *Pou2f3*^-/-^ SI epithelial monolayers stimulated as indicated. (**D**) Quantification of water content of fecal pellets collected from indicated mice immediately after oral treatment with C8 (30 mg/kg). In the graphs, each symbol represents an individual mouse pooled from two or more experiments. In (A-B) groups represent sequential stimulations of the same tissue and in (C) groups represent individual monolayers. *p < 0.05, **p < 0.01, ***p < 0.001, ****p < 0.001 by multiple Mann-Whitney tests with Holm Sídák’s multiple comparisons test (A, D), or multiple unpaired t tests with Holm Sídák’s multiple comparisons test (C). ns, not significant. Graphs depict mean +/-SEM.

### Tuft cell-derived ACh is not required for ILC2 activation

Having defined ACh-dependent fluid secretion as a tuft cell effector function in unmanipulated mice, we next considered the role of tuft cell-derived ACh during Type 2 immunity. Consistent with previous reports in the airways,^63^ we found that tamoxifen partially suppressed Type 2 responses in the intestine. For example, treating protist-colonized *Chat^fl/fl^;Pou2f3^Cre-Ert^*^2^*^/+^* mice with tamoxifen for one week reduced tuft cell counts by nearly half in both WT and Cre+ mice (Fig. S4A). The effect of tamoxifen was less noticeable during helminth infection, perhaps because this induces more Type 2 inflammation (Fig. S4B). Nonetheless, given these non-specific effects of tamoxifen, we focused on identifying *Chat*-dependent effects by analyzing *Chat*-sufficient and *Chat*-deficient mice that had all been treated with tamoxifen.

Previously characterized SI tuft cell effector molecules (e.g. IL-25 and LTC_4_) have primarily been shown to regulate ILC2 activation.^21, 27^ Also, recent studies have reported that ILC2s express *Chat* following activation and that ACh can potentiate their production of cytokines and proliferation, perhaps via autocrine signaling^64, 65^. We therefore asked if tuft cell-derived ACh was signaling to ILC2s in addition to inducing fluid secretion. We began by testing if ACh could enhance ILC2 activation using an *in vitro* model of acute (6-hour) ILC2 stimulation.^27^ Pairing ACh with IL-25 to mimic the results of tuft cell activation, we failed to detect any ACh-dependent ILC2 activation as measured by IL-13 reporter expression and secretion of IL-13 and IL-5 (Fig. S4C-D). By contrast, LTC_4_ greatly enhanced ILC2 activation when given with IL-25, as expected. We conclude that ACh does not induce cytokine expression in ILC2s sorted from unmanipulated mice and is therefore unlikely to contribute to their initial activation.

**Supplemental Figure 4:**
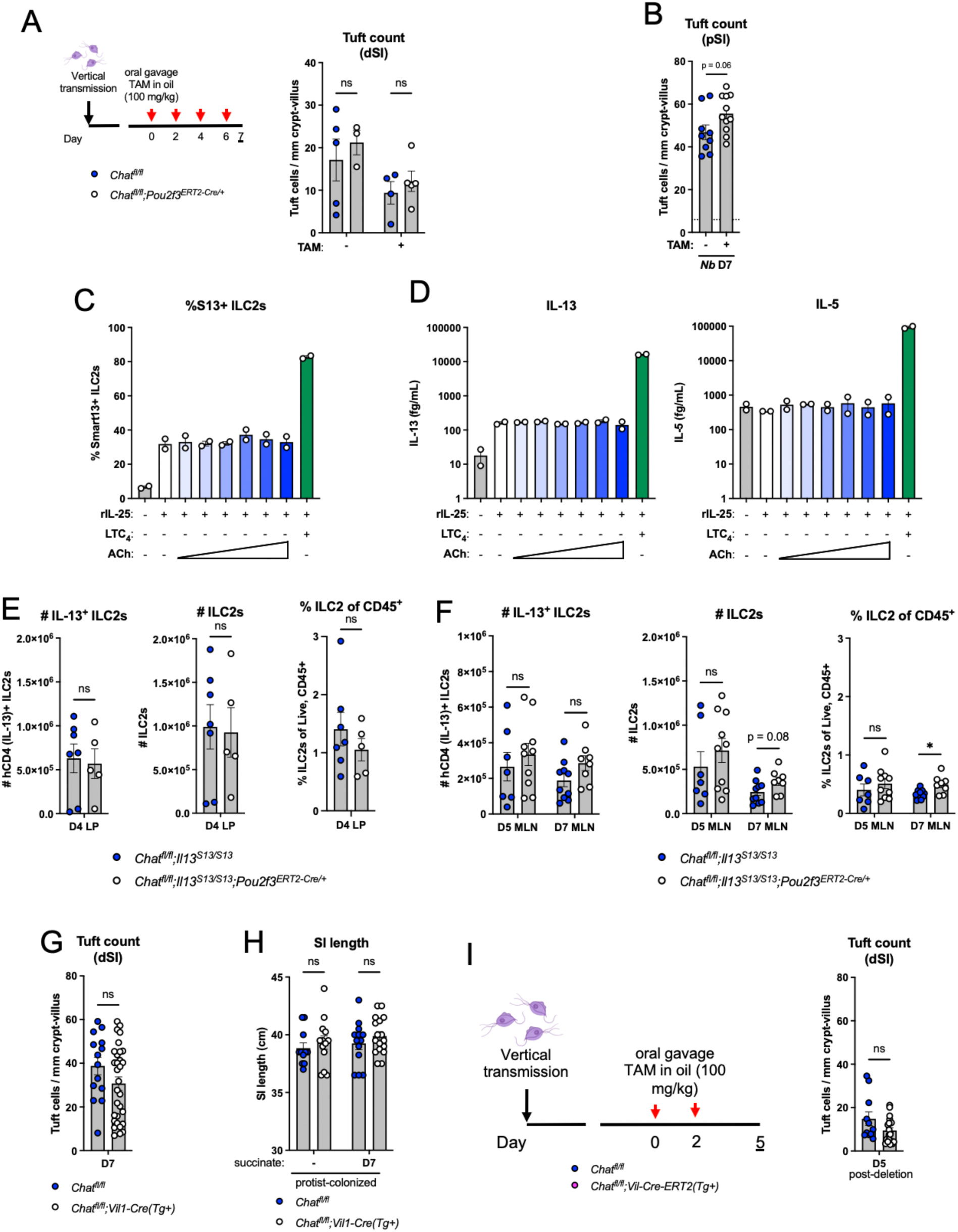
(**A**) Schematic of tamoxifen (TAM) treatment of protist-colonized *Chat-fl; Pou2f3^ERT^*^2^*^-Cre/+^* mice and quantification of dSI tuft counts by immunofluorescence at D7 after start of treatment. (**B**) Quantification of pSI tuft counts by immunofluorescence of WT mice treated with or without tamoxifen and infected with *Nb* for 7 days. (**C** and **D**) (C) Flow cytometric quantification of percent hCD4+ (IL-13+) SILP ILC2s and (D) IL-13 and IL-5 concentration in their supernatant after 6 hr *in vitro* culture with the indicated conditions (0.1 ng/mL rIL-25, serial 10-fold dilutions of ACh from 10 mM to 0.1 µM, 1 nM LTC4). (**E** and **F**) Quantification of number of hCD4+ (IL-13+) ILC2s, total ILC2s, and percent ILC2s at the indicated timepoints, tissues, and genotypes. (**G**) Quantification of dSI tuft counts by immunofluorescence from indicated mice treated with 150 mM succinate drinking water for 7 days. (**H**) SI length from indicated mice vertically-colonized with *T. rainier* protists with or without 7 days of additional 150 mM succinate drinking water treatment. (**I**) Schematic of acute deletion of *Chat* from vertically *T. musculis* (*Tm*) *-*colonized mice and quantification of dSI tuft counts by immunofluorescence 5 days after start of treatment. In the (A-B, E-I), each symbol represents an individual mouse from two or more pooled experiments. In (C and D) each symbol represents a technical replicate of cells sorted from pooled mice. *p < 0.05, **p < 0.01, ***p < 0.001, ****p < 0.001 by multiple Mann-Whitney tests with Holm Sídák’s multiple comparisons test (A, F, H) or Mann-Whitney test (B, E, G, I). ns, not significant. Graphs depict mean +/-SEM.

To test if tuft cell-derived ACh was involved in ILC2 activation *in vivo* and at later timepoints, we generated *Chat^fl/fl^;Pou2f3^ERT^*^2^*^-Cre/+^* mice that also expressed an IL-13 reporter (Smart13). We treated these mice with tamoxifen, infected with *Nb*, and assessed early ILC2 activation in the LP 4 days post infection (dpi), about 2 days after the worms arrive in the intestine. We found no difference in ILC2 activation or expansion (Fig. 4A, S4E). Since *Chat* has been detected in activated but not resting ILC2s, we also tested if tuft cell-derived ACh regulated ILC2s later during infection. Given the difficulty of isolating viable cells from Type 2 inflamed SI, we assessed ILC2s in the mesenteric lymph nodes at 5 and 7 dpi. Again, we saw equivalent activation and expansion of ILC2s between *Chat*-sufficient and -deficient mice (Fig. 4B, S4F). We conclude that tuft cell-derived ACh does not contribute to ILC2 activation during helminth infection of the SI.

**Figure 4:**
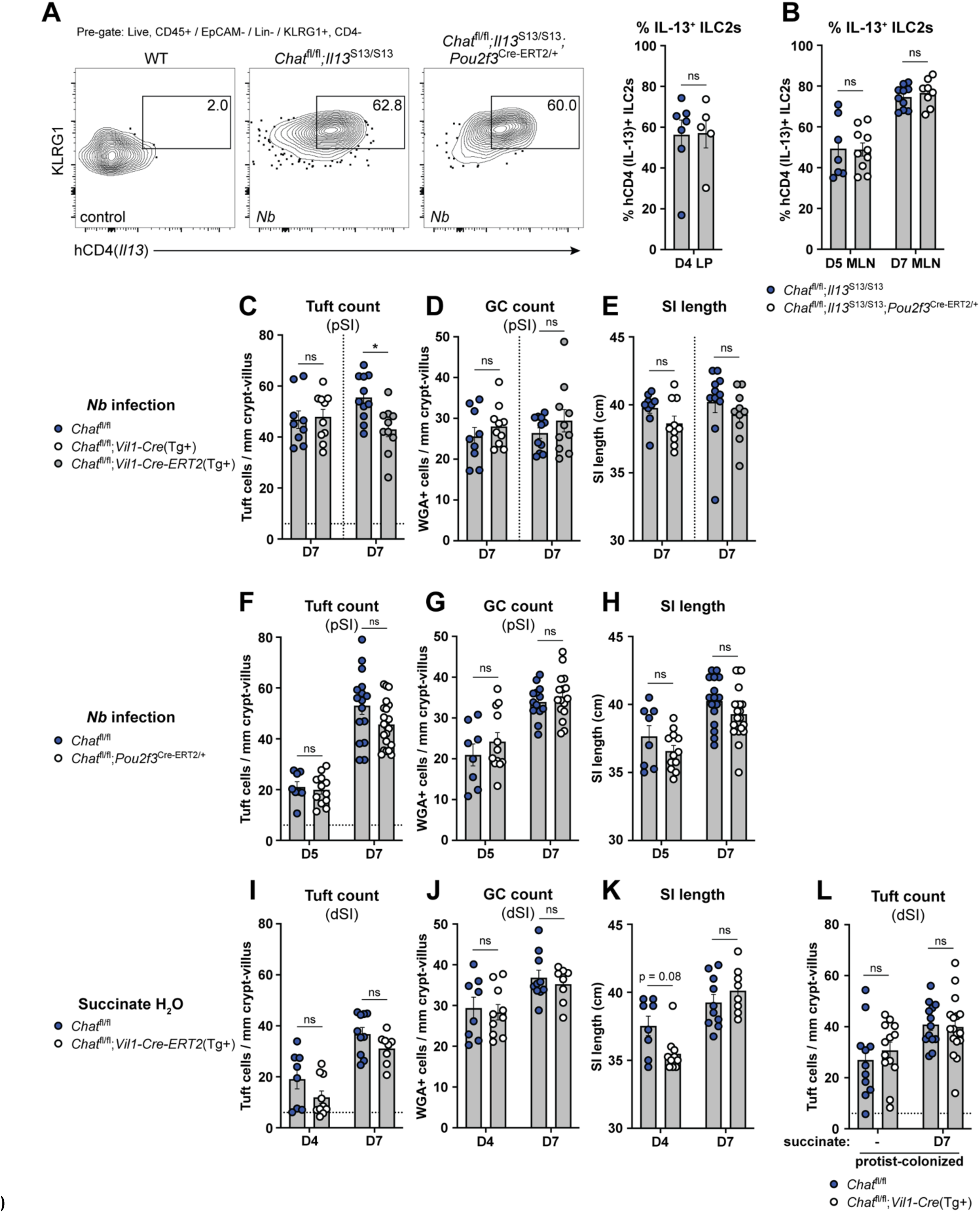
Tuft cell-derived ACh is not required for ILC2 activation or intestinal remodeling. (**A** and **B**) (A) Representative flow cytometry and quantification of percent hCD4^+^ (IL-13^+^) ILC2s (Lin^-^, CD45^+^, KLRG1^+^, CD4^-^) in the SI LP and (B) mesenteric lymph nodes (MLN) at the indicated *Nb* infection timepoints. (**C**, **D**, and **E**) (C) Quantification of pSI tuft cells (DCLK1+) and (D) goblet cells (WGA+) by immunofluorescence and (E) total SI length from the indicated mice at D7 of *Nb* infection. (**F**, **G**, and **H**) Same analysis as in C-E in the indicated mice at the indicated *Nb* infection timepoints. (**I**, **J**, and **K**) Same analysis as in C-E in the indicated mice at the indicated timepoints of 150 mM succinate drinking water treatment. (**L**) Quantification of tuft cells (DCLK1+) by immunofluorescence from indicated mice vertically-colonized with *T. rainier* protists with or without 7 days of additional 150 mM succinate drinking water treatment. In the graphs, each symbol represents an individual mouse from two or more pooled experiments. For graphs of tuft cell counts, horizontal dashed line signifies baseline tuft cell count in unmanipulated mice. *p < 0.05, **p < 0.01, ***p < 0.001, ****p < 0.001 by Mann Whitney test (A) or multiple Mann-Whitney tests with Holm Sídák’s multiple comparisons test (B-L). ns, not significant. Graphs depict mean +/-SEM. Also see Fig. S4.

### Tuft cell-derived ACh is not required for Type 2 intestinal remodeling

An effective immune response to helminths requires intestinal remodeling, such as hyperplasia of tuft cells and mucus-producing GCs and lengthening of the SI.^23, 66^ IL-13 is critical for this process, but ACh might also contribute. For example, ACh has been implicated in direct modulation of epithelial cell differentiation,^67, 68^ and could also impact tissue remodeling via other AChR-expressing cells, such as neurons.^55^

Using tuft cell frequency, goblet cell frequency, and intestinal length as markers of SI remodeling, we found little or no defect 7 days after *Nb* infection of mice with either constitutive (*Vil1-Cre(Tg+)*) or acute (*Vil1-Cre-Ert2(Tg+)*) deletion of *Chat* in IECs (Fig. 4C-E). The same was true both 5 and 7 days after *Nb* infection when we used *Pou2f3-Cre-ERT2* for tuft cell-specific *Chat* deletion (Fig. 4F-H). Since tuft cell circuits are distinct in the pSI and dSI,^11, 27^ we also tested Type 2 remodeling in the dSI 4 and 7 days after starting treatment with succinate-supplemented drinking water. As before, there was little effect of either constitutive or inducible IEC *Chat* deletion (Fig. 4I-K; S4G). Likewise, *Chat^fl/fl^;Vil1-Cre(Tg+)* mice vertically colonized with protists had no defect in tuft cell hyperplasia or SI length at homeostasis or following one week of additional succinate drinking water treatment (Fig. 4L, S4H). Acute loss of *Chat* in protist*-*colonized *Chat^fl/fl^;Vil1-Cre-ERT2(Tg+)* mice also had no effect on tuft cell hyperplasia 5 days later (Fig. S4I). In sum, Type 2 remodeling is broadly intact in the absence of tuft cell ACh. We did observe small but significant decreases in tuft cell hyperplasia or SI lengthening in some assays, but this effect was inconsistent and minimal compared to loss of other tuft cell effector molecules (e.g. IL-25 and LTC_4_).^21, 27^

### Tuft cell hyperplasia results in enhanced ACh-dependent fluid secretion

The ability of tuft cells to induce fluid secretion in the steady state led us to ask how it would change during Type 2 inflammation, when tuft cell numbers can increase 10-fold and fluid secretion might support worm clearance as part of the canonical weep and sweep response. First, we asked whether the number and frequency of *Chat*+ tuft cells changed with Type 2 inflammation. While the frequency of *Chat*+ tuft cells decreased (Fig. S5A), this was more than compensated for by the hyperplasia, such that the total number of *Chat+* tuft cells per millimeter crypt/villus was increased in the pSI and especially the dSI 7 days after either succinate-treatment or *Nb*-infection (Fig. 5A).

**Figure 5:**
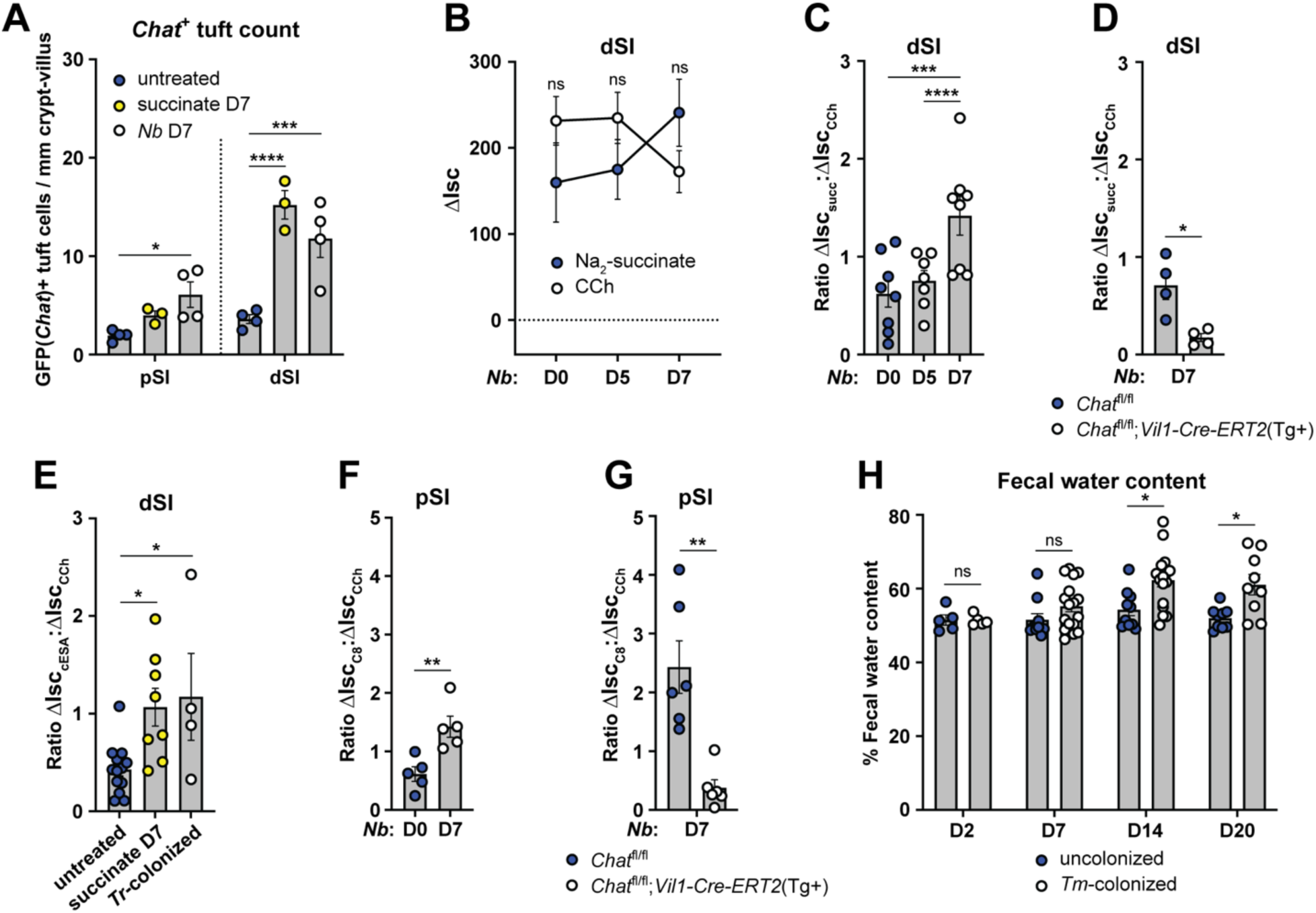
Tuft cell hyperplasia results in enhanced ACh-dependent fluid secretion. (**A**) Quantification of GFP(*Chat*)*+* tuft cells (*Il25-*RFP*+*) from the pSI and dSI of WT mice untreated, treated with 150 mM Na2-succinate drinking water (succinate), or infected with *N. brasiliensis* (*Nb*) for 7 days. (**B**) 1′Isc values of dSI from WT mice infected with *Nb* for the indicated number of days and stimulated as indicated (10 mM succinate, lumenal; 100 µM CCh, basolateral). (**C**) Ratio of succinate ΔIsc values to CCh 1′Isc values of dSI from (B). (**D**) Ratio of succinate ΔIsc values to CCh ΔIsc values of dSI of indicated mice 7 days after *Nb* infection. (**E**) Ratio of cESA ΔIsc values to CCh ΔIsc values of dSI from WT mice treated with succinate (as in A) or vertically colonized with *T. rainier* (*Tr*) protists. (**F** and **G**) Ratio of C8 1′Isc values to CCh ΔIsc values of pSI from (F) WT or (G) mice of indicated genotypes mice at indicated timepoints after *Nb* infection. (**H**) Quantification of water content of fecal pellets collected at indicated timepoints post *T. musculis* (*Tm*) colonization of WT mice. In the graphs, each symbol represents an individual mouse pooled from two or more experiments. *p < 0.05, **p < 0.01, ***p < 0.001, ****p < 0.001 by two way ANOVA with Dunnett’s multiple comparisons test (A), Mann-Whitney test (B, D, F-G), one way ANOVA (C, E), or multiple Mann-Whitney tests with Holm Sídák’s multiple comparisons test (H). ns, not significant. Graphs depict mean +/-SEM. Also see Fig. S5.

Next, we infected mice with *Nb* to induce tuft cell hyperplasia and quantified the fluid secretion response to succinate in the pSI and dSI across the course of infection. Although pSI tuft cells do not respond to succinate at steady state, we wondered if those that emerge during infection might be responsive. The pSI did not become responsive to succinate (or cESA) over the course of infection, but the dSI succinate response increased by day 7, when tuft cell numbers peaked (Fig. 5B, S5B-C). The increased response to succinate was less than the ∼3-fold increase of *Chat+* tuft cells that we observed on D7 of *Nb* infection, likely because the epithelium’s total capacity to respond to ACh/CCh was reduced by Type 2 inflammation, as previously reported (Fig. 5B).^69^

This effect could be quantified by measuring the ratio of succinate-induced 1′Isc to CCh-induced 1′Isc (Fig. 5C), highlighting how hyperplasia allows tuft cells to capture a greater proportion of the total epithelial ACh response. We confirmed that the increased succinate response on day 7 still required epithelium-derived ACh (Fig. 5D, S5D) and did not occur without *Sucnr1* (Fig. S5E). Enhanced succinate/cESA-induced fluid secretion was also observed if tuft cell hyperplasia was established with oral succinate or *Tritrichomonas* colonization (Fig. 5E). We found that tuft cell hyperplasia induced by *Nb* infection also increased C8-dependent fluid secretion in both the pSI and dSI, and that this required epithelial *Chat* (Fig. 5F-G, S5F-K).

To further test the hypothesis that increased numbers of tuft cells drive increased succinate-induced fluid secretion during Type 2 remodeling, we turned to Balb/c mice. Unmanipulated Balb/c mice are nearly tuft cell-deficient in the dSI, but activation of ILC2s with exogenous IL-25 increases tuft cell numbers (Fig. S5L).^44^ Accordingly, unmanipulated Balb/c dSI did not respond to cESA in the Ussing chamber, but responsiveness was induced by rIL-25 treatment, indicating that increased numbers of ACh-producing tuft cells were needed (Fig. S5M).

Lastly, to test if *in vivo* fluid secretion was enhanced during Type 2 remodeling, we measured the fecal water content of protist-colonized mice. Compared to uncolonized mice, mice colonized with the succinate-producing protist *T. musculis* had increased fecal water content by 14 and 20 days after colonization (Fig. 5H). This suggested that tuft cells were repeatedly responding to protist-derived succinate and inducing fluid secretion. Thus, the increased number of *Chat*+ tuft cells induced during SI Type 2 inflammation drives increased ACh-dependent fluid secretion despite an overall desensitization of the tissue to ACh.

**Supplemental Figure 5:**
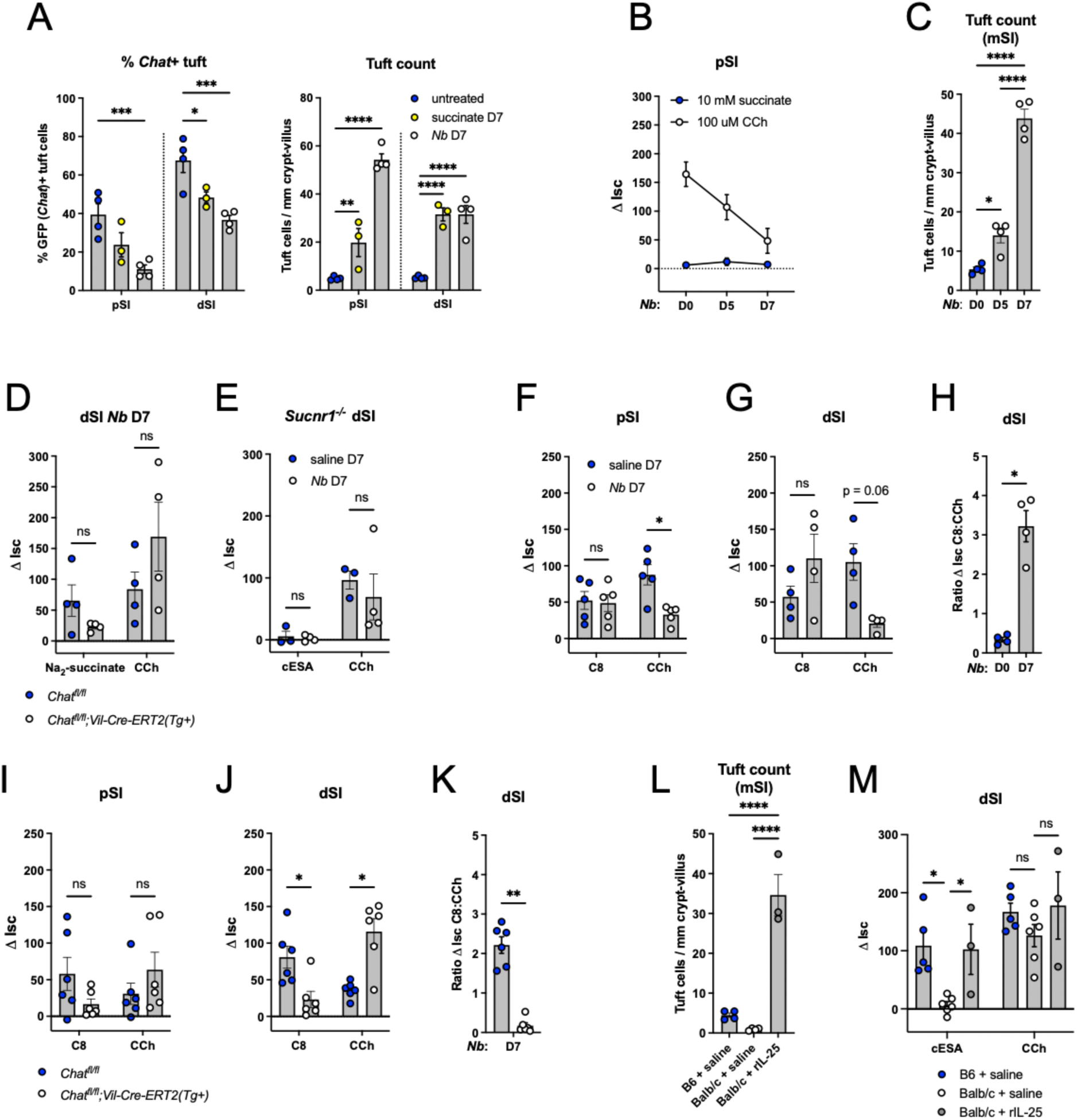
(**A**) Quantification of percent GFP(*Chat*)+ tuft cells and RFP(*Il25*)+ tuft cells from the pSI and dSI of WT mice untreated, treated with 150 mM Na2-succinate drinking water (succinate), or infected with *N. brasiliensis* (*Nb*) for the duration indicated. (**B**) 1′Isc values of pSI from WT mice infected with *Nb* for the indicated number of days and stimulated as indicated (10 mM succinate, lumenal; 100 µM CCh, basolateral). (**C**) Quantification of tuft cells (DCLK1+) by immunofluorescence from medial SI (mSI) of mice in (B). (**D**) 1′Isc values of dSI from mice of indicated genotypes infected with *Nb* for 7 days and stimulated as indicated. (**E**) 1′Isc values of *Sucnr1^-/-^* dSI with or without 7 day *Nb* infection, stimulated as indicated. (**F-G**) 1′Isc values of (F) pSI and (G) dSI from WT mice with or without 7 day *Nb* infection, stimulated as indicated. (**H**) Ratio of C8 ΔIsc values to CCh ΔIsc values in (G). (**I** and **J**) 1′Isc values of (I) pSI and (J) dSI from mice of indicated genotype infected with *Nb* for 7 days, stimulated as indicated. (**K**) Ratio of C8 ΔIsc values to CCh ΔIsc values from (J). (**L**) Quantification of tuft cells (DCLK1+) by immunofluorescence in the mSI of mice of indicated genotypes treated as indicated. (**M**) 1′Isc values of dSI from mice in (L) stimulated as indicated. In the graphs, each symbol represents an individual mouse (one tissue or average of two) from two or more pooled experiments. *p < 0.05, **p < 0.01, ***p < 0.001, ****p < 0.001 by two way ANOVA with Dunnett’s multiple comparisons test (A) two way ANOVA with Tukey’s multiple comparisons test (M), Mann-Whitney test (B, H, K), one way ANOVA with Tukey’s multiple comparisons test (C, L), or multiple Mann-Whitney tests with Holm Sídák’s multiple comparisons test (D-G, I-J). ns, not significant. Graphs depict mean +/-SEM.

### Tuft cell ACh regulates helminth clearance but not protist colonization

There is little evidence to suggest that tuft cell sensing of *Tritrichomonas sp.* and the resulting Type 2 immune response in the SI alter the total abundance of protists,^22^ but we wondered if tuft cell induced fluid secretion might instead regulate protist localization along the length of the SI, with the goal of containing protists to the dSI and cecum. We therefore assessed the abundance of protists across the pSI, dSI, and cecum of vertically-colonized *Chat^fl/fl^;Vil1-Cre* mice (Fig. S6A), hypothesizing an increase of protists in the pSI of *Chat*-deficient mice. Constitutive deletion of *Chat* in tuft cells had no effect on protist abundance or localization across the SI or cecum (Fig. S6B). Treating protist-colonized mice with succinate to amplify Type 2 immunity also did not uncover a phenotype (Fig. S6B), and acute deletion of *Chat* for 5 days likewise failed to alter protist abundance or localization (Fig. S6C). Thus, the physiologic function of protist sensing by tuft cells remains unclear.

**Supplemental Figure 6:**
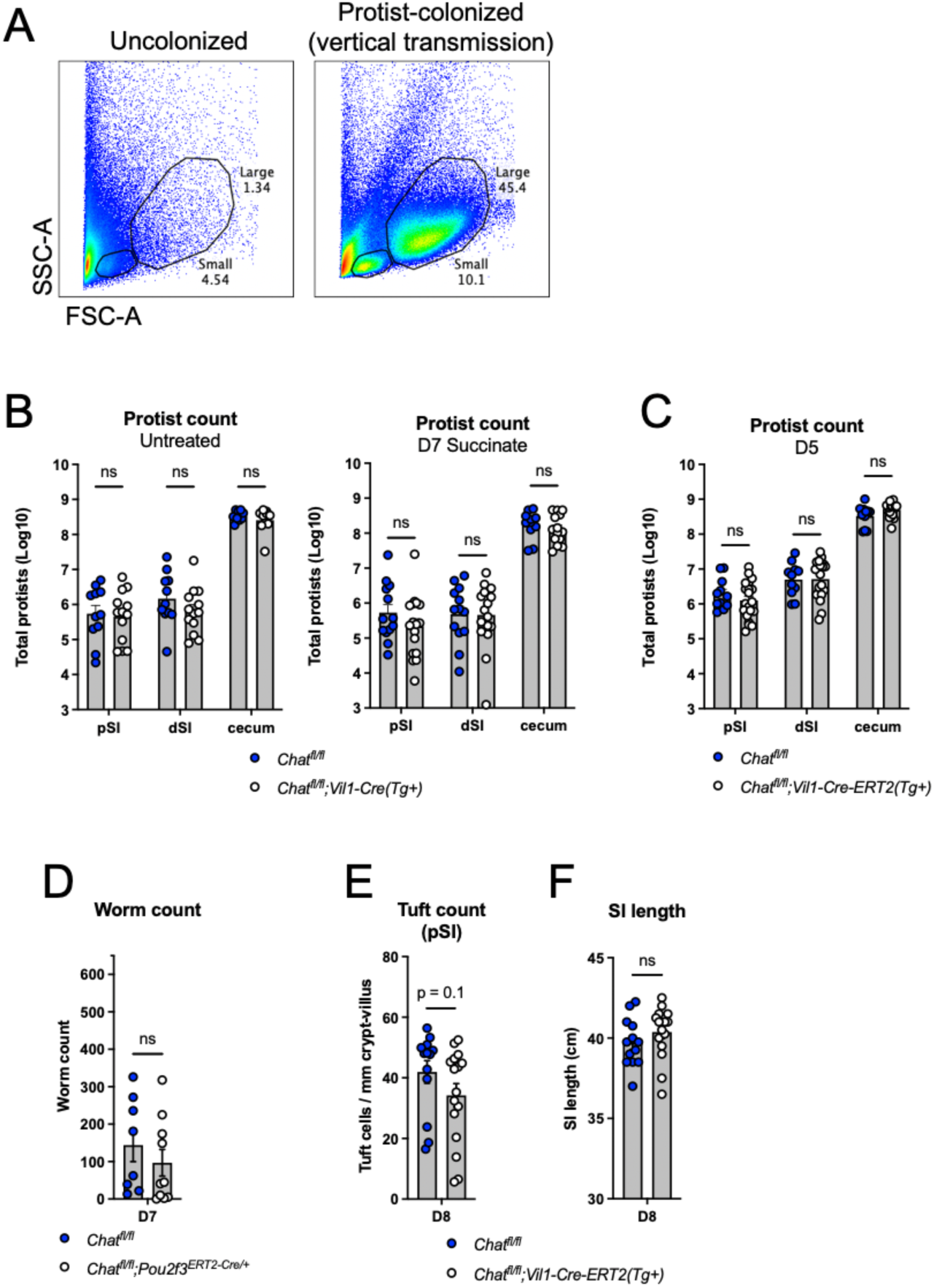
(**A**) Representative flow cytometry of cecal contents from uncolonized and protist-colonized mice showing gating of protists by size. The “Large” gate contains *Tritrichomonas sp.* protists. (**B**) Quantification of total protists by flow cytometry in indicated tissues of vertically-colonized mice of indicated genotypes left untreated or given 150 mM Na2-succinate in drinking water for 7 days. (**C**) Quantification of total protists by flow cytometry in indicated tissues of vertically-colonized mice of indicated genotypes administered tamoxifen 5 days prior to analysis. (**D**) Quantification of total SI *Nb* in mice of indicated genotype infected with *Nb* for 7 days without tamoxifen administration. (**E** and **F**) (E) Quantification of tuft cells (DCLK1+) by immunofluorescence and (F) total SI length 8 days post *Nb* infection of mice of indicated genotype given a single dose of tamoxifen (125 mg/kg) on day 5. In the graphs, each symbol represents an individual mouse (one tissue or average of two) from two or more pooled experiments. *p < 0.05, **p < 0.01, ***p < 0.001, ****p < 0.001 by multiple Mann-Whitney tests with Holm Sídák’s multiple comparisons test (B-C) or Mann-Whitney test (D-F). ns, not significant. Graphs depict mean +/-SEM.

On the other hand, the requirement for tuft cell sensing and downstream Type 2 immunity for clearing helminths from the SI is well established.^11, 20, 45^ To test if tuft cell ACh contributes to helminth clearance, despite not impacting ILC2 activation or tissue remodeling, we assessed worm burden in mice lacking epithelial *Chat* at 7 dpi, a timepoint when WT mice begin to clear worms from the SI. Indeed, both constitutive and acute deletion of epithelial *Chat* led to an increased SI worm burden (Fig. 6A). Tuft-specific *Chat* deletion using *Pou2f3-Cre-Ert2* led to the same clearance delay 7 dpi (Fig. 6B). Worm burdens were equivalent in CRE-positive and -negative mice 5 dpi, suggesting normal colonization of the SI by *Nb* arriving from the lung. The delayed clearance was also not due to the loss of one *Pou2f3* allele in *Chat^fl/fl^;Pou2f3^Cre-ERT^*^2^*^/+^* mice as they cleared worms normally when not treated with tamoxifen (Fig. S6D). In order allow initiation of type 2 remodeling to proceed normally and delete tuft cell ACh only during worm clearance, we waited until 5 dpi to administer a single dose of tamoxifen to *Chat^fl/fl^;Vil1-Cre-ERT2* mice. Consistent with our earlier observation that tamoxifen suppresses intestinal Type 2 immunity, worm clearance in WT mice was delayed to day 8, but we again found a *Chat-*dependent delay in worm clearance despite normal intestinal remodeling. (Fig. 6C, Fig. S6E-F). Thus, we propose that tuft-cell derived ACh contributes to worm clearance by the induction of epithelial fluid secretion.

**Figure 6:**
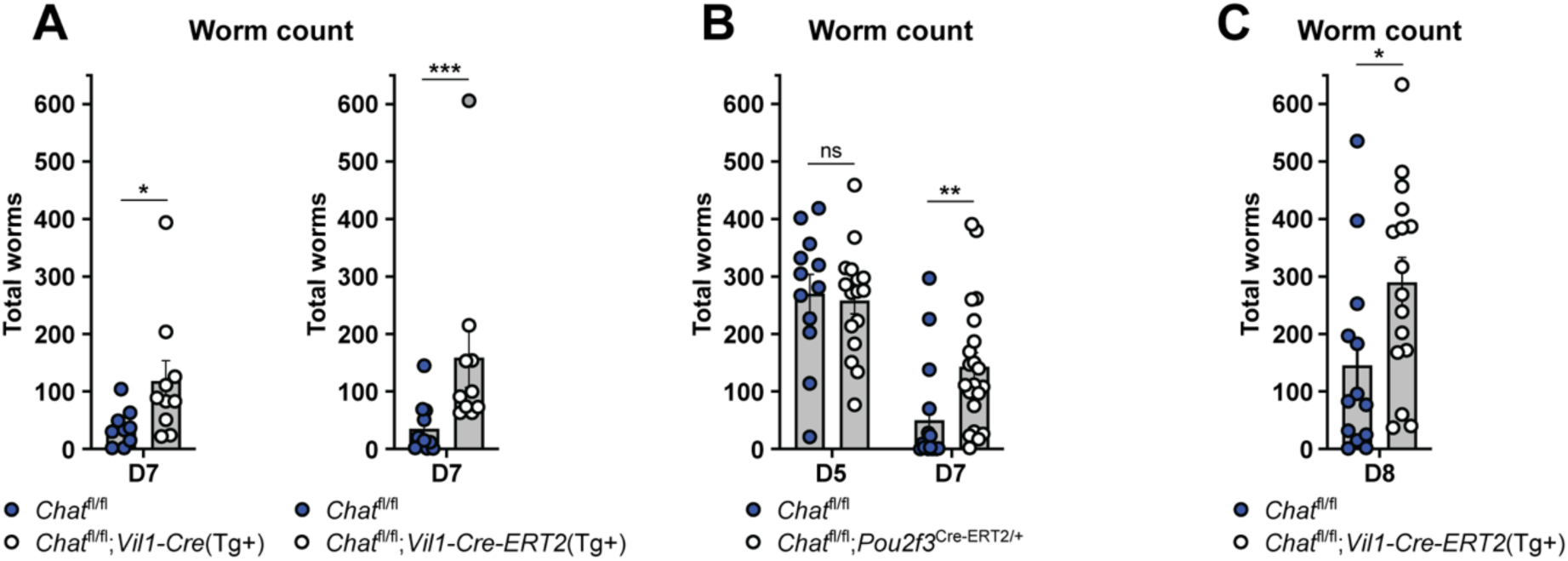
Tuft cell-derived ACh contributes to helminth clearance. (**A** and **B**) Quantification of total SI *Nb* in mice of indicated genotypes at (A) 7 or (B) indicated days post infection. (**C**) Quantification of total SI *Nb* 8 days post infection in mice of indicated genotypes given single dose of tamoxifen on D5. In the graphs, each symbol represents an individual mouse pooled from two or more experiments. *p < 0.05, **p < 0.01, ***p < 0.001, ****p < 0.001 by Mann-Whitney test (A, C), or multiple Mann-Whitney tests with Holm Sídák’s multiple comparisons test (B). ns, not significant. Graphs depict mean +/-SEM. Also see Fig. S6.

## Discussion

This study identifies an epithelium-intrinsic response unit that couples tuft cell chemosensing to epithelial fluid secretion via the release of ACh from tuft cells. This effector function is common to tuft cells in multiple tissues and is executed within seconds of activation. In the SI, tuft cell-derived ACh is required for timely clearance of helminth infection, but unlike all previously identified tuft cell effector functions, tuft cell-derived ACh does not impact ILC2 activation nor downstream intestinal remodeling. Instead, it appears that tuft cell ACh provides an acute signal that contributes to the weep and sweep responses that push worms out of the intestine. The magnitude of tuft cell-dependent fluid secretion correlates with the number of *Chat*+ tuft cells, suggesting one possible function for the tuft cell hyperplasia that occurs even after initial sensing of the helminth has been achieved.

Some details of tuft cell-regulated fluid secretion remain unresolved. For example, we do not understand the regulation of *Chat* in SI tuft cells. It is unclear why only a subset of SI tuft cells are *Chat*+, why there is a proximal to distal gradient of *Chat+* tuft cells, and why the frequency of *Chat+* tuft cells decreases during Type 2 inflammation (although the total number increases due to tuft cell hyperplasia). Furthermore, most tuft cells do not express *Slc5a7*, which encodes CHT1, the transporter that neurons use to import choline for ACh synthesis, nor *Slc18a3*, which encodes VAChT, a transporter that loads ACh into secretory vesicles in neurons.^70^ Lastly, it remains to be seen whether tuft cell-derived ACh also induces bicarbonate secretion, as this often occurs together with chloride release and further supports the unfolding of extracellular mucus.^71^

We have also not identified the precise ACh receptor(s) that mediate(s) the fluid secretion response, although inhibition by atropine implicates a muscarinic rather than nicotinic receptor. *Chrm1* and *Chrm3* are the only detectable muscarinic receptor transcripts in unmanipulated SI epithelium^9, 11^ and *Nb* clearance is delayed in *Chrm3*^-/-^ mice.^40^ Induction of Type 2 cytokines is also impaired in these *Chrm3^-/-^* mice, so conditional *Chrm* alleles will be needed to identify in which cells the receptors are required and for which aspects of Type 2 immunity.

While tuft cell-derived ACh was not required for ILC2 activation during helminth infection and intestinal remodeling was largely intact, there were slight yet significant defects in tuft cell numbers or SI length at some timepoints analyzed. Prior literature has shown that deletion of *Chrm3* from the SI epithelium causes an *increase* in tuft cell numbers at baseline but a *decrease* in tuft cells following irradiation.^67^ Perhaps epithelial ACh signaling has damage-induced functions that overlap with helminth-induced intestinal remodeling. Regulation of intestinal epithelial differentiation by tuft cell-derived ACh bears further study.

We have focused on tuft cell ACh signaling on enterocytes to drive fluid secretion, but ACh receptors are expressed by many cells, including other types of intestinal epithelial cells. Goblet cells undergo compound exocytosis of mucus in response to ACh,^46, 47, 72^ and the formation of goblet-associated antigen passages (GAPs) has also been linked to ACh signaling.^73, 74^ Additionally, tuft cell ACh was recently reported to induce mucus secretion from cholangiocytes in the gallbladder.^57^ We therefore extensively tested the hypothesis that ACh from SI tuft cells signals on villus goblet cells to induce mucus secretion and GAP formation, but we could not find any evidence that this occurs *in vivo* (data not shown). A recent study demonstrating that muscarinic receptor expression is restricted to GCs at the base of SI crypts, and that only these cells respond acutely to CCh,^75^ may explain why we did not detect tuft cell-dependent regulation of GC mucus secretion. While it remains possible that ACh from SI crypt tuft cells regulates secretion by GCs (and Paneth cells), we generally found fewer *Chat*+ tuft cells in the crypts than in the villi, and chemosensing pathways are likely not yet functional in immature crypt tuft cells. Perhaps in the colon tuft cell ACh regulates the function of sentinel GCs at crypt openings.^76^ Lastly, ACh-regulated smooth muscle contraction is critical for SI helminth clearance^28, 37^ and tuft cells have been linked to smooth muscle function in other tissues.^19, 57^ The short extracellular half-life of ACh combined with the distance between epithelial tuft cells and smooth muscles that surround the SI make direct signaling unlikely. Nonetheless, while we did not see evidence of direct contact between tuft cells and enteric neurons, they have been previously reported in the SI^10, 77, 78^ and thus we cannot rule out the possibility that tuft cell ACh regulates smooth muscle contraction via the enteric nervous system.

During Type 2 inflammation in the SI, the maximal fluid secretion induced by ACh is dramatically reduced,^69^ possibly to prevent excessive fluid loss or diarrhea during chronic helminth infection. At the same time, the number of *Chat*+ tuft cells increases, such that tuft cell-regulated fluid secretion is maintained or even enhanced compared to baseline. This re-wiring of ACh-regulated fluid secretion may represent a regulatory mechanism that minimizes fluid loss due to endogenous signals while maintaining the ability to respond to lumen-restricted agonists such as helminths via tuft cell sensing. In that regard, tuft cells also have an advantage over mast cells and neurons, which can induce enhanced fluid secretion during Type 2 inflammation via release of histamine and/or prostaglandin E2,^69^ but can only respond to ligands that penetrate the mucosal barrier. Relatedly, it remains unclear whether the acetylcholinesterases secreted by helminths can penetrate the mucosal barrier to target ACh in the tissue, or whether helminths are only able to counter the effects of ACh during tissue-dwelling phases of their lifecycle.

Although we have focused on the SI in this study, we propose tuft cells link chemosensing to fluid secretion in all tissues. Indeed, with the exception of tuft-ILC2 circuit activation, all other known tuft cell effector functions occur instantaneously and seem to mediate evasion (e.g. breathing cessation)^14, 16^ and expulsion (e.g. mucociliary sweep) of microbes and other agonists.^8, 17^ Fluid secretion fits this paradigm. The mucosal barrier must be constantly hydrated and fluid secretion can provide a flushing effect. Based on the ligands tuft cells sense in different tissues, such mechanisms could be important to clear allergens from the upper airways or bacteria from the trachea and urethra. Tuft cell sensing may also reduce baseline fluid secretion, as one recent study suggested.^79^ The ligands and function of tuft cells in the colon are only just being elucidated,^56^ but we predict that tuft cell-regulated fluid secretion would help maintain a healthy mucosal barrier here too. Tuft cell frequency is generally decreased in patients with active inflammatory bowel disease,^80, 81^ consistent with a role for tuft cells in preventing bacterial infiltration. Conversely, increased tuft cell frequency was detected in colonic biopsies from patients with diarrhea-predominant irritable bowel syndrome, a largely non-inflammatory condition of unknown origin.^82^

Why tuft cells sense succinate in either the SI or, as we have now demonstrated, the trachea, remains unclear. We could not find any impact of tuft cell *Chat* deletion on *Tritrichomonas* burden or distribution, but perhaps tuft cell-mediated fluid secretion, together with IL-13-induced anti-microbial peptides and mucus production, acts more locally to keep microbes away from the epithelium.^83^ Succinate levels have also been shown to increase in contexts of bacterial dysbiosis, and inducing tuft cell hyperplasia with succinate treatment reduces inflammation in a model of ileitis.^80^ As for the trachea, aberrant release of cellular succinate into the airways, which occurs in some patients with cystic fibrosis, can promote colonization and biofilm formation by the pathosymbiont *Pseudomonas aeruginosa.*^84^ Induction of CFTR-dependent fluid secretion by prostaglandin E2 released from airway tuft cells has also been suggested.^86^ Thus, tracheal tuft cells may induce fluid secretion to flush away succinate and other soluble molecules and to deter bacterial accumulation.

Therapeutically, benefit may be achieved by tuning tuft cell effector functions up or down, depending on the context and need. For example, tuft cell-induced fluid secretion may prove useful in treating cystic fibrosis patients in whom CFTR-dependent fluid secretion is impaired, while certain patients suffering from diarrhea might benefit from reduced tuft cell function. Future study should investigate the involvement of tuft cell-induced fluid secretion in human disease.

## Acknowledgements

We thank all members of the von Moltke lab for helpful discussion and input on this manuscript. We thank D. Hailey and the Garvey Cell Imaging Lab in the Institute for Stem Cell & Regenerative Medicine for microscopy support; the mouse husbandry staff in the UW SLU vivarium; V. Gersuk, K. O’Brien and the Benaroya Research Institute Genomics Core for help with RNA sequencing; and M. F. Fontana for helpful comments on the manuscript. Flow cytometry data were acquired through the University of Washington, Cell Analysis Facility Shared Resource Lab, with NIH award 1S10OD024979-01A1 funding for the Symphony A3. We thank Takeda for the generous gift of the Class 8 TRPM5 agonist.

## Contributions

TEB designed and performed experiments, analyzed data, and wrote the paper with JVM. LMW, DBS, MMM, DNK, and JWM assisted with experiments at the University of Washington. CF and MBG performed *in vivo* Class 8 and *T. musculis* fecal water content experiments at Stanford University with supervision and funding provided by MRH. KAB and LMR performed pilot Ussing chamber experiments with supervision and funding provided by JSD. AB, MS, and RM led the development and validation of the Class 8 TRPM5 agonist at Takeda Pharmaceuticals in collaboration with Evotec. JVM conceived of and supervised the study, analyzed data, acquired funding, and wrote the paper with TEB.

## Methods

### Study Design

All experiments were performed using randomly assigned mice without investigator blinding. No data were excluded, except from Ussing chambers when mounted tissues failed to respond to stimulation by a positive control (e.g. CCh). All data points reflect biological replicates, except for *in vitro* ILC2 stimulations and epithelial monolayer experiments where each data point is a technical replicate. Data were pooled from multiple experiments unless otherwise noted. The number of independent experiments is included in the figure legends.

### Experimental Animals

Mice aged 6 weeks and older were used for all experiments. Mice were age-matched within each experiment. Pooled results include both male and female mice of varying ages unless otherwise indicated. Mouse strains used in this study are listed in Table S3. Acute deletion of conditional alleles in mice was achieved by oral gavage with tamoxifen dissolved in corn oil (100 mg/kg). *Chat^fl/fl^;Vil1-Cre-ERT2(Tg+)* mice were administered tamoxifen on day -4 and 0 of infection with *N. brasiliensis* and on day -6 and -4 of treatment with 150 mM succinate drinking water, or as noted in the text.

*Chat^fl/fl^;Pou2f3^Cre-ERT^*^2^*^/+^* mice were administered tamoxifen every other day starting on day -4 of *N. brasiliensis* infection or as noted in the text. *DREADD(Tg+); Pou2f3^Cre-ERT^*^2^*^/+^* mice were administered tamoxifen chow for 7 days prior to Ussing experiments. All mice (CRE+ and CRE-) received tamoxifen treatment. All mice were maintained in specific pathogen-free conditions at the University of Washington or Stanford University and were confirmed to be free of *Tritrichomonas sp.* by microscopy and qPCR, unless specifically colonized for experimental purposes. All procedures were conducted within University of Washington or Stanford University (Class 8 gavage) IACUC guidelines under approved protocols.

### Measuring epithelial ion flux with Ussing chambers

Ussing chamber protocols were informed by Clarke et al.^51^

*Tissue mounting*: Mice 7-12 weeks of age were euthanized by CO_2_ and segments of the intestine (SI, cecum, colon) were harvested and flushed with cold Krebs Buffered Ringer’s solution + mannitol (10 mM D-mannitol, 115 mM NaCl, 2.4 mM K_2_HPO_4_, 0.4 mM KH_2_PO_4_, 25 mM NaHCO_3_, 1.2 mM CaCl_2_ dihydrate, 1.2 MgCl_2_ hexahydrate; pH 7.25-7.4). Intestines (first 5 cm of pSI and last 5 cm of dSI) were fileted open along the mesenteric line, trimmed and mounted to the pins of an Ussing chamber cassette with aperture of 0.3 cm^2^ (Physiologic Instruments, Reno, USA), avoiding Peyer’s patches. For some experiments the SI was pinned to a Sylgard-coated plate with the serosal side up and the muscle layer scored with a scalpel and “stripped” away using forceps under a dissection scope. The resulting epithelium-submucosa tissue was mounted in a cassette as normal. For trachea preparations, the entire trachea was harvested, surrounding tissue (esophagus) removed, and cut open lengthwise along the anterior side (away from esophagus) for mounting in a 2 mm^2^ aperture cassette (Physiologic Instruments, Reno, USA). Cassettes containing tissues were mounted in Ussing chambers (Physiologic Instruments, Reno, USA) and the lumenal chambers filled with 5 mL of KBR + mannitol and the basolateral chambers filled with 5 mL of KBR + glucose (10 mM D-glucose, 115 mM NaCl, 2.4 mM K_2_HPO_4_, 0.4 mM KH_2_PO_4_, 25 mM NaHCO_3_, 1.2 mM CaCl_2_ dihydrate, 1.2 MgCl_2_ hexahydrate; pH 7.25-7.4). The chambers were warmed to 37C and bubbled with carbogen (95% O_2_, 5% CO_2_) for the duration of the experiment. For Cl^-^ replacement experiments, Cl^-^ free KBR was used, in which gluconate is substituted for Cl-(115 mM D-gluconic acid sodium salt, 2.4 mM K_2_HPO_4_, 0.4 mM KH_2_PO_4_, 25 mM NaHCO_3_, 4 mM calcium D-gluconate monohydrate, 1.2 mM magnesium D-gluconate hydrate; pH 7.25-7.4).

*Measuring the short circuit current (Isc)*: Automatic voltage clamping was performed by MultiChannel Voltage-Current Clamp (Physiologic Instruments, Reno, USA). Voltage differences between electrodes and fluid resistance of the buffer were compensated prior to insertion of the tissue cassette. Isc was measured by voltage clamp every 1 second and recorded using Acquire & Analyze software (Physiologic Instruments, Reno, USA) and normalized to tissue area. After a 20 min equilibration period, tissues were stimulated lumenally with Na_2_-succinate (1, 10 mM) or cis-epoxysuccinic acid (10 mM), basolaterally with 5 µM Class 8, or bilaterally with 1 µM Compound 21. For succinate/cESA stimulation, subsequent stimulations were administered every 10 min; for Class 8 the interval was increased to 15 min due to the slower kinetics of the response. Subsequent stimulations included lumenal 20 mM NaCl, basolateral 100 µM CCh, and bilateral cocktail of 100 µM 3-Isobutyl-1-methylxanthine (IBMX) + 10 µM forskolin. Chemical inhibitors were administered 5 min after start of equilibration period (15 min before first stimulation): basolateral 100 µM bumetanide, bilateral 10 µM ibuprofen, basolateral 10 µM atropine, basolateral 1 µM tetrodotoxin, and lumenal 1 mM carbenoxolone. ΔIsc values were calculated as the difference between the Isc measurement at the peak of the stim response and the Isc measurement taken right before adding the agonist. For tissues that did not respond to the agonist (e.g., knockout mouse intestines, tissues treated with inhibitors) and therefore had no peak response, the Isc value was taken at the same timepoint as the peak Isc value for the corresponding WT or control tissue.

### Measuring fecal water content

Mice were orally gavaged with vehicle (0.5% methylcellulose + 1% Tween 20) or 30 mg/kg Class 8, and fecal samples (2+ pellets) were taken at 0 and 3 hours post gavage. For protist-colonized experiments, B6 mice were colonized with *T. musculis* protists and fecal samples collected at 2, 7,14, and 20 days post colonization. Fecal samples were dried at 60C overnight and % water content calculated as (1-(dry weight/wet weight))*100.

### Monolayer culture and cysteinyl leukotriene ELISA

Proximal small intestine was isolated and villi were gently scraped off with a glass coverslip. Tissue was incubated for 30 minutes at 4° with 2mM EDTA to release epithelial crypts, then washed twice with cold PBS and filtered through a 70 um strainer. Crypts were resuspended in complete monolayer media (DMEM/F12 supplemented with 2mM glutamine, 100U/mL penicillin, 100mg/mL streptomycin, 10mM HEPES, N2 supplement, B27 supplement, R-spondin (10% supernatants from R-spondin secreting cells), Noggin (10% supernatant from Noggin secreting cells), 500mM N-acetylcysteine, 50ug/mL mEGF, and 10 µM Y27632). Plates were coated with 2% Matrigel in cold DMEM/F12 and incubated at 37° for at least 30 minutes. Media was aspirated from the plate, and 1000 crypts were plated per well of a 48-well plate. Crypts were incubated overnight, and non-adherent cells were aspirated the next day. Test stimuli diluted in HBSS containing Ca2+/Mg2+ were added to monolayers and stimulated at 37° for 30 minutes. Supernatants were collected and used for the Cysteinyl Leukotriene Express ELISA kit (Cayman Chemical) according to manufacturer’s protocol.

### Succinate and cytokine treatment

For succinate experiments mice were given 150mM sodium succinate hexahydrate (Thermo) ad libitum in drinking water for the indicated amount of time. Recombinant murine IL-25 (500 ng; R&D) was given for 3 consecutive days intraperitoneally in 200 µL PBS.

### Helminth infections and analysis

*N. brasiliensis* larvae were raised and maintained as previously described.^85^

Mice were infected subcutaneously with 500 *N. brasiliensis* L3. At sacrifice, the entire SI was fileted open and total worms counted under a stereomicroscope.

### Protist colonization and analysis

Breeding pairs were colonized with *Tritrichomonas musculis* or *T. rainier* as previously described.^87^ Pups from colonized breeding pairs were analyzed. Protist numbers were quantified by flow cytometry as described by Chudnovskiy et al.^88^ Briefly, 10 cm of pSI and dSI were flushed into a 15 mL conical with 10 mL RT PBS using a gavage needle. Cecal contents were harvested into 15 mL conical, weighed, and then 10 mL RT PBS added. Samples were let sit for 30 min at RT, vortexed, and then passed through a 70 µm filter. Protists were washed, stained with DAPI, then count beads added and data collected on a FACSCanto II (BD Biosciences). Protist numbers per gram of cecal content was calculated.

### Intestinal tissue fixation and staining

Intestinal tissues were flushed with PBS and fixed in 4% paraformaldehyde for 3-4 hours at 4°C, washed with PBS, and incubated in 30% (w/v) sucrose overnight at 4°C.

Samples were then coiled into “Swiss rolls”, embedded in Optimal Cutting Temperature Compound (Tissue-Tek) and sectioned at 8 μm on a CM1950 cryostat (Leica).

Immunofluorescent staining was performed in PBS with 1% BSA at room temperature as follows: 1 hr 10% donkey serum with 1:1000 Fc Block, 1 hr (or O/N at 4°C) primary antibody, 5 min wash, 45 min secondary donkey antibody and/or WGA-488, 5 min wash, and mounted with Vectashield plus DAPI (Vector Laboratories). Images were acquired with an Axio Observer A1 (Zeiss) microscope with a 10X or 20X A Plan objective. Tuft cell frequency was calculated using ImageJ software to manually quantify DCLK1^+^ cells per millimeter of crypt-villus axis. Goblet cell frequency was calculated using ImageJ software to manually quantify total WGA+ cells in the villus (crypts were excluded because WGA also labels Paneth cells) per millimeter of crypt villus axis. For each replicate, four 10x images of the Swiss roll were analyzed and at least 25 total villi counted.

### Single-cell tissue preparation for flow cytometry

For single cell epithelial preparations from SI, tissues were flushed with PBS, Peyer’s patches removed, opened longitudinally, and rinsed with PBS. Tissue was cut into small pieces, shaken vigorously for 20 seconds in 30 mL cold HBSS (Ca^+2^/Mg^+2^-free) with 1 mM HEPES, drained, and then incubated rocking at 37°C for 10 min in 15 mL HBSS (Ca^+2^/Mg^+2^-free) supplemented with 3 mM EDTA and 1 mM HEPES. Tissues were vortexed thoroughly and released epithelial cells passed through a 70 μm filter.

This process was repeated for a total of 3 rounds. Supernatants were pooled and washed with HBSS (Ca^+2^/Mg^+2^-free) with 1 mM HEPES before staining for flow cytometry.

For lamina propria (LP) preparations from uninfected mice, SI was processed as above to remove the epithelial fraction. Tissues were then incubated shaking at 37°C for 30 minutes in 10 mL RPMI 1640 supplemented with 20% FCS, 1 mM HEPES, 0.05 mg/ml DNase I (Sigma Aldrich), and 1 mg/mL Collagenase A (Sigma Aldrich). Tissues were vortexed and cells were passed through a 100 μm filter, then a 40 μm filter, washing with cold HBSS (Ca^+2^/Mg^+2^-free) with 1 mM HEPES. Cells were washed and stained for flow cytometry.

For LP preparations from *N. brasiliensis*-infected mice (D4), mice were anaesthetized with 5% avertin. The peritoneal cavity was opened, the SI nicked at the junction with the stomach and transected at the cecum and flushed with 20 mL of 37°C HBSS (Ca^+2^/Mg^+2^-free) plus 1 mM HEPES. Then the mice were perfused through the heart with 30 mL of 37°C HBSS (Ca^+2^/Mg^+2^-free) with 30 mM EDTA and 1 mM HEPES. Three minutes after perfusion was completed, the first 10 cm of the proximal SI was harvested, Peyer’s patches removed, opened longitudinally, and cut into small pieces and shaken vigorously for 20 seconds in 30 mL cold HBSS (Ca^+2^/Mg^+2^-free) with 1 mM HEPES, then drained. Tissues were then digested and processed as above in uninfected mice.

For mesenteric lymph node (MLN) preparations, SI-draining MLN were harvested into RPMI + 5% FBS on ice, mashed through a 70 μm filter, the filter washed with RPMI + 5% FBS, and cells washed and stained for flow cytometry.

### Flow cytometry and cell sorting

Single cell suspensions from tissues were prepared as described above. For flow cytometry, SI epithelium and MLN samples were stained in DPBS (Ca^+2^/Mg^+2^-free) with 3% FCS and LP samples were stained in PBS (Ca^+2^/Mg^+2^-free) with 3% FCS, 2 mM EDTA, and 0.02 mg/mL DNase I with antibodies to surface markers for 30 min at 4°C, followed by DAPI (Roche) for dead cell exclusion. When cell counts were needed, counting beads (Spherotech) were added prior to running flow cytometry. Samples were run on a FACSCanto II or LSRII (BD Biosciences) and analyzed with FlowJo 10.8.1.

Samples were FSC-A/SSC-A gated to exclude debris, FSC-A/FSC-H gated to select single cells, and gated to exclude dead cells. For cell sorting, single cell suspensions were prepared and stained as described and sorted on an Aria III (BD Biosciences).

### ILC2 Stimulation Assay

Entire SILP from several mice were pooled and ILC2s (EpCAM-, CD45^+^, Lin(CD3, CD4, CD5, CD8, CD11b, CD19, NK1.1, FcER1)^-^, KLRG1^+^) sorted as described. Sorted cells were plated at 5000 cells per well in a 96-well plate and incubated at 37°C overnight in 10 ng/ml IL-7 (R&D Systems) and basal media composed of high glucose DMEM supplemented with non-essential amino acids, 10% FBS, 100 U/mL penicillin, 100mg/mL streptomycin, 10mM HEPES, 1mM sodium pyruvate, 100μM 2-mercaptoethanol, and 2mM L-glutamine. The next morning, media was replaced and cells were stimulated with the indicated agonist. After a six-hour stimulation, supernatant was collected and the cells were washed and stained with 1 uL/well of PE-conjugated anti-human CD4 for 20 min at 4°C. Cells were washed, resuspended in DAPI and analyzed on a CantoRUO (BD Biosciences). Cytokine levels in supernatants were measured using Enhanced Sensitivity Flex Sets (BD Biosciences) for mouse IL-5 and IL-13 according to the manufacturer’s protocol. Data was collected on an LSRII (BD Biosciences).

### RNA Sequencing and Analysis

150-200 tuft cells were sorted directly into lysis buffer from the SMART-Seq v4 Ultra Low Input RNA Kit (Takara) and cDNA generated following manufacturer’s instructions. Cells were sorted from four individual mice for each experiment. Sequencing libraries were generated using the Nextera XT library preparation kit with multiplexing primers, according to manufacturer’s protocol (Illumina), and library quality assessed using Tapestation (Agilent). High throughput sequencing was performed on NextSeq 2000 (Illumina), sequencing dual-indexed and paired-end 59 base pair reads. All samples were in the same run with a target depth of 5 million reads. Base calls were processed to FASTQs on BaseSpace (Illumina), and a base call quality-trimming step was applied to remove low-confidence base calls from the ends of reads. The FASTQs were aligned to the GRCm38 mouse reference genome, using STAR v.2.4.2a and gene counts were generated using htseq-count. Further analysis of the data was performed using the DIY.Transcriptomics (diytranscriptomics.com) pipeline, with experiment-specific modifications. Samples were filtered to exclude genes with counts per million = 0 in 4 or more samples and genes annotated as pseudogenes. Finally, samples were normalized to each other. To identify differentially expressed genes, precision weights were first applied to each gene based on its mean-variance relationship using VOOM,^89^ then data was normalized using the TMM method^90^ in EdgeR.^91^ Linear modeling and bayesian stats were employed via Limma^92^ to find genes that were up-or down-regulated by 2-fold (Log2FC = 1) or more, with a false-discovery rate (FDR) of 0.01. The code and results for these analyses are included as Data File S1 and S2.

### Statistical Analysis

Statistical analysis was performed as noted in figure legends using Prism 9 (GraphPad) software. Graphs show mean +/-SEM.

**Table S3.**
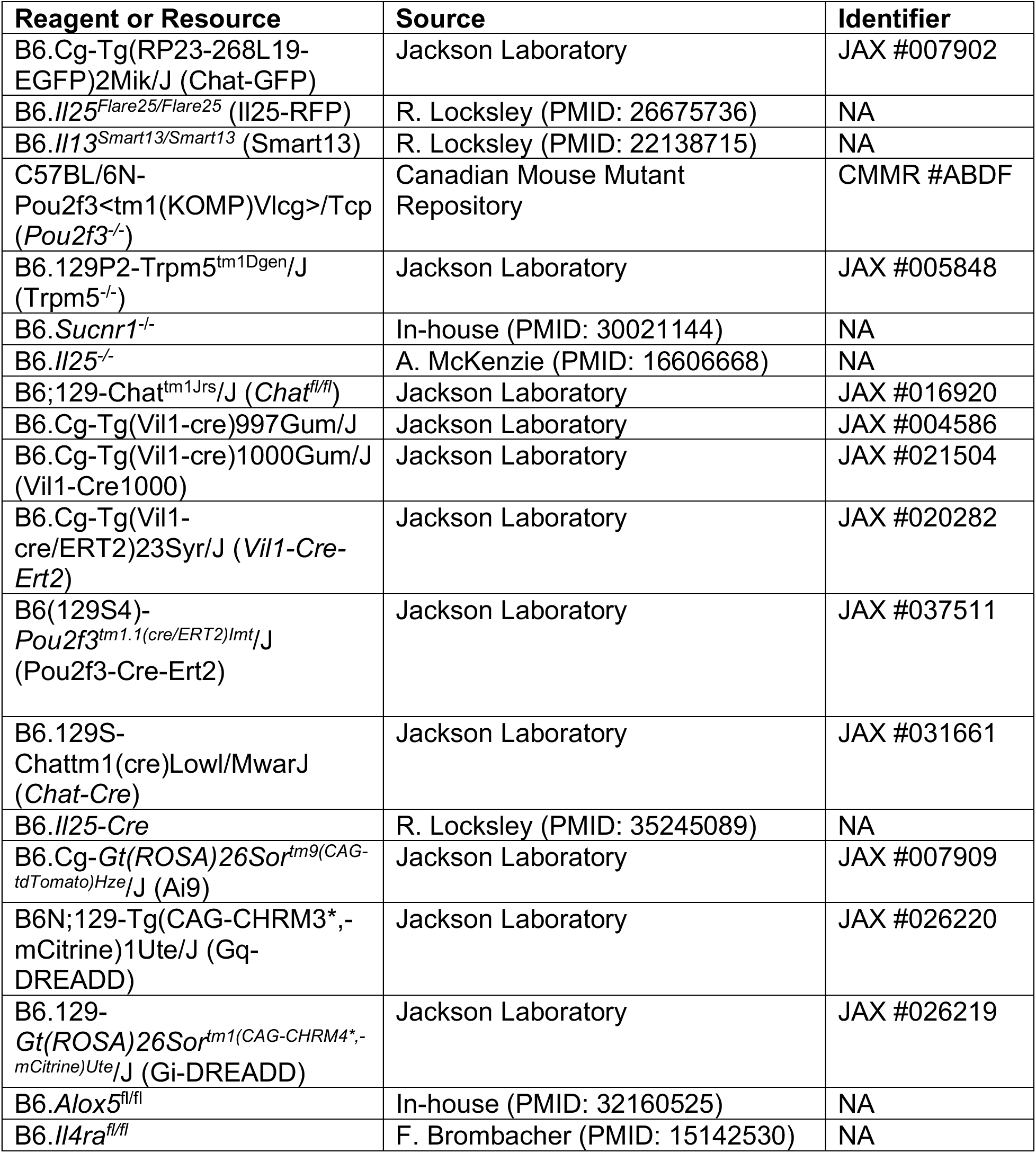

**Table S4.**

| **Reagent or Resource** | **Dilution Factor** | **Source** | **Identifier** |
| --- | --- | --- | --- |
| Rabbit α-DCLK1 | 1:1000 | Abcam | Cat#ab31704 |
| Rabbit α-TUJ1 (Beta-III tubulin) | 1:500 | Abcam | Cat#ab18207 |
| Goat α-GFP | 1:500 | Novus Bio | Cat#NB100-1770 |
| Rabbit α-dsRed | 1:500 | Clontech | Cat#632496 |
| Rabbit α-HA, clone 16B12 | 1:1000 | Biolegend | Cat#901516 |
| WGA-488 | 1:150 | Thermo | Cat#W11261 |
| Donkey α-Rabbit IgG AF594 | 1:1000 | Thermo | Cat#A-21207 |
| Donkey α-Goat IgG AF488 | 1:500 | Thermo | Cat#A-11055 |
| CD16 / CD32, clone 2.4G2 | 1:1000 | Tonbo | Cat# 70-0161-M001 |
| CD3 PerCP-Cy5.5, clone 145-2C11 | 1:100 | Biolegend | Cat#100328 |
| CD3 BV421, clone 145-2C11 | 1:400 | Biolegend | Cat#100335 |
| CD4 BV711, clone RM4-5 | 1:250 | Biolegend | Cat#100549 |
| CD4 eF450, clone RM4-5 | 1:200 | eBioscience | Cat# 48-0042-80 |
| hCD4 PE, clone RPA-T4 | 1:50 | Biolegend | Cat#300508 |
| CD5 PerCP-Cy5.5, clone 53-7.3 | 1:500 | Biolegend | Cat#100624 |
| CD5 eF450, clone 53-7.3 | 1:400 | Biolegend | Cat#100607 |
| CD8 PerCP-Cy5.5, clone 53-6.7 | 1:200 | Biolegend | Cat#100724 |
| CD8 BV421, clone 53-6.7 | 1:400 | Biolegend | Cat#100737 |
| CD11b AF700, clone M1/70 | 1:250 | Biolegend | Cat#101222 |
| CD11b BV421, clone M1/70 | 1:400 | Biolegend | Cat# 101235 |
| CD19 PerCP-Cy5.5, clone 6D5 | 1:250 | Biolegend | Cat#115533 |
| CD19 BV421, clone 6D5 | 1:400 | Biolegend | Cat# 115537 |
| CD24 PE, clone M1/69 | 1:300 | Biolegend | Cat#101807 |
| CD24 PerCP-Cy5.5, clone M1/69 | 1:300 | Biolegend | Cat#101824 |
| CD45 BV605, clone 30F11 | 1:300 | Biolegend | Cat#103155 |
| CD45 BV650, clone 30F11 | 1:500 | Biolegend | Cat#103151 |
| EpCAM PE-Dazzle, clone G8.8 | 1:300 | Biolegend | Cat#118235 |
| EpCAM AF488, clone G8.8 | 1:300 | Biolegend | Cat#118210 |
| EpCAM PE-Cy7, clone G8.8 | 1:300 | Biolegend | Cat#118215 |
| FcER1 BV421, clone Mar-1 | 1:400 | Biolegend | Cat#334623 |
| IL17RB APC, clone 9B10 | 1:100 | Biolegend | Cat#146307 |
| KLRG1 PE-Cy7, clone 2F1 | 1:250 | Biolegend | Cat#138416 |
| NK1.1 PerCP-Cy5.5, clone PK136 | 1:100 | Biolegend | Cat#108728 |
| NK1.1 BV421, clone PK136 | 1:200 | Biolegend | Cat#108731 |
| Siglec-F APC-Cy7, clone E50-2440 | 1:100 | BD | Cat#565527 |
| Siglec-F AF647, clone E50-2440 | 1:100 | BD | Cat# 562680 |
| Thy1.2 (CD90.2) BV605, clone 53-2.1 | 1:500 | Biolegend | Cat#140318 |

## References

1. Frizzell, R.A., and Hanrahan, J.W. (2012). Physiology of epithelial chloride and fluid secretion. Cold Spring Harb Perspect Med 2, a009563. 10.1101/cshperspect.a009563.

2. Cooke, H.J. (1998). “Enteric Tears”: Chloride Secretion and Its Neural Regulation. News Physiol Sci 13, 269–274.

3. Xue, J., Askwith, C., Javed, N.H., and Cooke, H.J. (2007). Autonomic nervous system and secretion across the intestinal mucosal surface. Auton Neurosci 133, 55–63. 10.1016/j.autneu.2007.02.001.

4. Hirota, C.L., and McKay, D.M. (2006). Cholinergic regulation of epithelial ion transport in the mammalian intestine. Br J Pharmacol 149, 463–479. 10.1038/sj.bjp.0706889.

5. Cox, M.A., Bassi, C., Saunders, M.E., Nechanitzky, R., Morgado-Palacin, I., Zheng, C., and Mak, T.W. (2020). Beyond neurotransmission: acetylcholine in immunity and inflammation. J Intern Med 287, 120–133. 10.1111/joim.13006.

6. Mashimo, M., Moriwaki, Y., Misawa, H., Kawashima, K., and Fujii, T. (2021). Regulation of Immune Functions by Non-Neuronal Acetylcholine (ACh) via Muscarinic and Nicotinic ACh Receptors. Int J Mol Sci 22, 6818. 10.3390/ijms22136818.

7. Yajima, T., Inoue, R., Matsumoto, M., and Yajima, M. (2011). Non-neuronal release of ACh plays a key role in secretory response to luminal propionate in rat colon. J Physiol 589, 953–962. 10.1113/jphysiol.2010.199976.

8. Perniss, A., Liu, S., Boonen, B., Keshavarz, M., Ruppert, A.-L., Timm, T., Pfeil, U., Soultanova, A., Kusumakshi, S., Delventhal, L., et al. (2020). Chemosensory Cell-Derived Acetylcholine Drives Tracheal Mucociliary Clearance in Response to Virulence-Associated Formyl Peptides. Immunity 52, 683–699.e11. 10.1016/j.immuni.2020.03.005.

9. Haber, A.L., Biton, M., Rogel, N., Herbst, R.H., Shekhar, K., Smillie, C., Burgin, G., Delorey, T.M., Howitt, M.R., Katz, Y., et al. (2017). A single-cell survey of the small intestinal epithelium. Nature 551, 333–339. 10.1038/nature24489.

10. Bezençon, C., Fürholz, A., Raymond, F., Mansourian, R., Métairon, S., Le Coutre, J., and Damak, S. (2008). Murine intestinal cells expressing Trpm5 are mostly brush cells and express markers of neuronal and inflammatory cells. J Comp Neurol 509, 514–525. 10.1002/cne.21768.

11. Nadjsombati, M.S., McGinty, J.W., Lyons-Cohen, M.R., Jaffe, J.B., DiPeso, L., Schneider, C., Miller, C.N., Pollack, J.L., Nagana Gowda, G.A., Fontana, M.F., et al. (2018). Detection of Succinate by Intestinal Tuft Cells Triggers a Type 2 Innate Immune Circuit. Immunity 49, 33–41.e7. 10.1016/j.immuni.2018.06.016.

12. Schütz, B., Ruppert, A.-L., Strobel, O., Lazarus, M., Urade, Y., Büchler, M.W., and Weihe, E. (2019). Distribution pattern and molecular signature of cholinergic tuft cells in human gastro-intestinal and pancreatic-biliary tract. Sci Rep 9, 17466. 10.1038/s41598-019-53997-3.

13. Deng, J., Tan, L.H., Kohanski, M.A., Kennedy, D.W., Bosso, J.V., Adappa, N.D., Palmer, J.N., Shi, J., and Cohen, N.A. (2021). Solitary chemosensory cells are innervated by trigeminal nerve endings and autoregulated by cholinergic receptors. Int Forum Allergy Rhinol 11, 877–884. 10.1002/alr.22695.

14. Tizzano, M., Gulbransen, B.D., Vandenbeuch, A., Clapp, T.R., Herman, J.P., Sibhatu, H.M., Churchill, M.E.A., Silver, W.L., Kinnamon, S.C., and Finger, T.E. (2010). Nasal chemosensory cells use bitter taste signaling to detect irritants and bacterial signals. Proc Natl Acad Sci U S A 107, 3210–3215. 10.1073/pnas.0911934107.

15. Saunders, C.J., Christensen, M., Finger, T.E., and Tizzano, M. (2014). Cholinergic neurotransmission links solitary chemosensory cells to nasal inflammation. Proc Natl Acad Sci U S A 111, 6075–6080. 10.1073/pnas.1402251111.

16. Krasteva, G., Canning, B.J., Hartmann, P., Veres, T.Z., Papadakis, T., Mühlfeld, C., Schliecker, K., Tallini, Y.N., Braun, A., Hackstein, H., et al. (2011). Cholinergic chemosensory cells in the trachea regulate breathing. Proc Natl Acad Sci U S A 108, 9478–9483. 10.1073/pnas.1019418108.

17. Hollenhorst, M.I., Jurastow, I., Nandigama, R., Appenzeller, S., Li, L., Vogel, J., Wiederhold, S., Althaus, M., Empting, M., Altmüller, J., et al. (2020). Tracheal brush cells release acetylcholine in response to bitter tastants for paracrine and autocrine signaling. FASEB J 34, 316–332. 10.1096/fj.201901314RR.

18. Lips, K.S., Wunsch, J., Zarghooni, S., Bschleipfer, T., Schukowski, K., Weidner, W., Wessler, I., Schwantes, U., Koepsell, H., and Kummer, W. (2007). Acetylcholine and molecular components of its synthesis and release machinery in the urothelium. Eur Urol 51, 1042–1053. 10.1016/j.eururo.2006.10.028.

19. Deckmann, K., Filipski, K., Krasteva-Christ, G., Fronius, M., Althaus, M., Rafiq, A., Papadakis, T., Renno, L., Jurastow, I., Wessels, L., et al. (2014). Bitter triggers acetylcholine release from polymodal urethral chemosensory cells and bladder reflexes. Proc Natl Acad Sci U S A 111, 8287–8292. 10.1073/pnas.1402436111.

20. Gerbe, F., Sidot, E., Smyth, D.J., Ohmoto, M., Matsumoto, I., Dardalhon, V., Cesses, P., Garnier, L., Pouzolles, M., Brulin, B., et al. (2016). Intestinal epithelial tuft cells initiate type 2 mucosal immunity to helminth parasites. Nature 529, 226–230. 10.1038/nature16527.

21. von Moltke, J., Ji, M., Liang, H.-E., and Locksley, R.M. (2016). Tuft-cell-derived IL-25 regulates an intestinal ILC2-epithelial response circuit. Nature 529, 221–225. 10.1038/nature16161.

22. Howitt, M.R., Lavoie, S., Michaud, M., Blum, A.M., Tran, S.V., Weinstock, J.V., Gallini, C.A., Redding, K., Margolskee, R.F., Osborne, L.C., et al. (2016). Tuft cells, taste-chemosensory cells, orchestrate parasite type 2 immunity in the gut. Science 351, 1329–1333. 10.1126/science.aaf1648.

23. Schneider, C., O’Leary, C.E., von Moltke, J., Liang, H.-E., Ang, Q.Y., Turnbaugh, P.J., Radhakrishnan, S., Pellizzon, M., Ma, A., and Locksley, R.M. (2018). A Metabolite-Triggered Tuft Cell-ILC2 Circuit Drives Small Intestinal Remodeling. Cell 174, 271–284.e14. 10.1016/j.cell.2018.05.014.

24. Lei, W., Ren, W., Ohmoto, M., Urban, J.F., Matsumoto, I., Margolskee, R.F., and Jiang, P. (2018). Activation of intestinal tuft cell-expressed Sucnr1 triggers type 2 immunity in the mouse small intestine. Proc Natl Acad Sci U S A 115, 5552–5557. 10.1073/pnas.1720758115.

25. Hofmann, T., Chubanov, V., Gudermann, T., and Montell, C. (2003). TRPM5 is a voltage-modulated and Ca(2+)-activated monovalent selective cation channel. Curr Biol 13, 1153–1158. 10.1016/s0960-9822(03)00431-7.

26. O’Leary, C.E., Schneider, C., and Locksley, R.M. (2019). Tuft Cells-Systemically Dispersed Sensory Epithelia Integrating Immune and Neural Circuitry. Annu Rev Immunol 37, 47–72. 10.1146/annurev-immunol-042718-041505.

27. McGinty, J.W., Ting, H.-A., Billipp, T.E., Nadjsombati, M.S., Khan, D.M., Barrett, N.A., Liang, H.-E., Matsumoto, I., and von Moltke, J. (2020). Tuft-Cell-Derived Leukotrienes Drive Rapid Anti-helminth Immunity in the Small Intestine but Are Dispensable for Anti-protist Immunity. Immunity 52, 528–541.e7. 10.1016/j.immuni.2020.02.005.

28. Marillier, R.G., Michels, C., Smith, E.M., Fick, L.C.E., Leeto, M., Dewals, B., Horsnell, W.G.C., and Brombacher, F. (2008). IL-4/IL-13 independent goblet cell hyperplasia in experimental helminth infections. BMC Immunol 9, 11. 10.1186/1471-2172-9-11.

29. McKenzie, G.J., Bancroft, A., Grencis, R.K., and McKenzie, A.N. (1998). A distinct role for interleukin-13 in Th2-cell-mediated immune responses. Curr Biol 8, 339–342. 10.1016/s0960-9822(98)70134-4.

30. Miller, H.R., Huntley, J.F., and Wallace, G.R. (1981). Immune exclusion and mucus trapping during the rapid expulsion of Nippostrongylus brasiliensis from primed rats. Immunology 44, 419–429.

31. Grencis, R.K. (2015). Immunity to Helminths: Resistance, Regulation, and Susceptibility to Gastrointestinal Nematodes. Annual Review of Immunology 33, 201–225. 10.1146/annurev-immunol-032713-120218.

32. Oeser, K., Schwartz, C., and Voehringer, D. (2015). Conditional IL-4/IL-13-deficient mice reveal a critical role of innate immune cells for protective immunity against gastrointestinal helminths. Mucosal Immunol 8, 672–682. 10.1038/mi.2014.101.

33. Herbert, D.R., Yang, J.-Q., Hogan, S.P., Groschwitz, K., Khodoun, M., Munitz, A., Orekov, T., Perkins, C., Wang, Q., Brombacher, F., et al. (2009). Intestinal epithelial cell secretion of RELM-beta protects against gastrointestinal worm infection. J Exp Med 206, 2947–2957. 10.1084/jem.20091268.

34. Hu, Z., Zhang, C., Sifuentes-Dominguez, L., Zarek, C.M., Propheter, D.C., Kuang, Z., Wang, Y., Pendse, M., Ruhn, K.A., Hassell, B., et al. (2021). Small proline-rich protein 2A is a gut bactericidal protein deployed during helminth infection. Science (New York, N.Y.) 374, eabe6723. 10.1126/science.abe6723.

35. Wu, D., Ahrens, R., Osterfeld, H., Noah, T.K., Groschwitz, K., Foster, P.S., Steinbrecher, K.A., Rothenberg, M.E., Shroyer, N.F., Matthaei, K.I., et al. (2011). Interleukin-13 (IL-13)/IL-13 receptor alpha1 (IL-13Ralpha1) signaling regulates intestinal epithelial cystic fibrosis transmembrane conductance regulator channel-dependent Cl-secretion. J Biol Chem 286, 13357–13369. 10.1074/jbc.M110.214965.

36. Horsnell, W.G.C., Cutler, A.J., Hoving, J.C., Mearns, H., Myburgh, E., Arendse, B., Finkelman, F.D., Owens, G.K., Erle, D., and Brombacher, F. (2007). Delayed goblet cell hyperplasia, acetylcholine receptor expression, and worm expulsion in SMC-specific IL-4Ralpha-deficient mice. PLoS Pathog 3, e1. 10.1371/journal.ppat.0030001.

37. Akiho, H., Blennerhassett, P., Deng, Y., and Collins, S.M. (2002). Role of IL-4, IL-13, and STAT6 in inflammation-induced hypercontractility of murine smooth muscle cells. Am J Physiol Gastrointest Liver Physiol 282, G226–232. 10.1152/ajpgi.2002.282.2.G226.

38. Zhao, A., McDermott, J., Urban, J.F., Gause, W., Madden, K.B., Yeung, K.A., Morris, S.C., Finkelman, F.D., and Shea-Donohue, T. (2003). Dependence of IL-4, IL-13, and nematode-induced alterations in murine small intestinal smooth muscle contractility on Stat6 and enteric nerves. J Immunol 171, 948–954. 10.4049/jimmunol.171.2.948.

39. Darby, M., Schnoeller, C., Vira, A., Culley, F.J., Bobat, S., Logan, E., Kirstein, F., Wess, J., Cunningham, A.F., Brombacher, F., et al. (2015). The M3 muscarinic receptor is required for optimal adaptive immunity to helminth and bacterial infection. PLoS Pathog 11, e1004636. 10.1371/journal.ppat.1004636.

40. McLean, L.P., Smith, A., Cheung, L., Urban, J.F., Sun, R., Grinchuk, V., Desai, N., Zhao, A., Raufman, J.-P., and Shea-Donohue, T. (2016). Type 3 muscarinic receptors contribute to intestinal mucosal homeostasis and clearance of Nippostrongylus brasiliensis through induction of TH2 cytokines. Am J Physiol Gastrointest Liver Physiol 311, G130–141. 10.1152/ajpgi.00461.2014.

41. Sanderson, B.E., and Ogilvie, B.M. (1971). A study of acetylcholinesterase throughout the life cycle of Nippostrongylus brasiliensis. Parasitology 62, 367–373. 10.1017/s0031182000077519.

42. Blackburn, C.C., and Selkirk, M.E. (1992). Characterisation of the secretory acetylcholinesterases from adult Nippostrongylus brasiliensis. Mol Biochem Parasitol 53, 79–88. 10.1016/0166-6851(92)90009-9.

43. Lawrence, C.E., and Pritchard, D.I. (1993). Differential secretion of acetylcholinesterase and proteases during the development of Heligmosomoides polygyrus. Int J Parasitol 23, 309–314. 10.1016/0020-7519(93)90004-i.

44. Nadjsombati, M.S., Niepoth, N., Webeck, L.M., Kennedy, E.A., Jones, D.L., Baldridge, M.T., Bendesky, A., and Moltke, J. von (2022). Genetic mapping reveals Pou2af2-dependent tuning of tuft cell differentiation and intestinal type 2 immunity. 2022.10.19.512785. 10.1101/2022.10.19.512785.

45. von Moltke, J., Ji, M., Liang, H.-E., and Locksley, R.M. (2016). Tuft-cell-derived IL-25 regulates an intestinal ILC2-epithelial response circuit. Nature 529, 221–225. 10.1038/nature16161.

46. Specian, R.D., and Neutra, M.R. (1980). Mechanism of rapid mucus secretion in goblet cells stimulated by acetylcholine. J Cell Biol 85, 626–640. 10.1083/jcb.85.3.626.

47. Gustafsson, J.K., Ermund, A., Johansson, M.E.V., Schütte, A., Hansson, G.C., and Sjövall, H. (2012). An ex vivo method for studying mucus formation, properties, and thickness in human colonic biopsies and mouse small and large intestinal explants. Am J Physiol Gastrointest Liver Physiol 302, G430–438. 10.1152/ajpgi.00405.2011.

48. Stockinger, S., Albers, T., Duerr, C.U., Ménard, S., Pütsep, K., Andersson, M., and Hornef, M.W. (2014). Interleukin-13-mediated paneth cell degranulation and antimicrobial peptide release. J Innate Immun 6, 530–541. 10.1159/000357644.

49. Hubel, K.A. (1976). Intestinal ion transport: effect of norepinephrine, pilocarpine, and atropine. Am J Physiol 231, 252–257. 10.1152/ajplegacy.1976.231.1.252.

50. Banks, M.R., and Farthing, M.J.G. (2002). Fluid and electrolyte transport in the small intestine. Curr Opin Gastroenterol 18, 176–181. 10.1097/00001574-200203000-00004.

51. Clarke, L.L. (2009). A guide to Ussing chamber studies of mouse intestine. Am J Physiol Gastrointest Liver Physiol 296, G1151–1166. 10.1152/ajpgi.90649.2008.

52. Browning, J.G., Hardcastle, J., Hardcastle, P.T., and Redfern, J.S. (1978). Localization of the effect of acetylcholine in regulating intestinal ion transport. J Physiol 281, 15–27. 10.1113/jphysiol.1978.sp012406.

53. Geubelle, P., Gilissen, J., Dilly, S., Poma, L., Dupuis, N., Laschet, C., Abboud, D., Inoue, A., Jouret, F., Pirotte, B., et al. (2017). Identification and pharmacological characterization of succinate receptor agonists. Br J Pharmacol 174, 796–808. 10.1111/bph.13738.

54. Vanoye, C.G., Altenberg, G.A., and Reuss, L. (1999). Inhibition of P-glycoprotein-mediated transport by a hydrophobic contaminant in commercial gluconate salts. Am J Physiol 276, C1439–1442. 10.1152/ajpcell.1999.276.6.C1439.

55. Harrington, A.M., Hutson, J.M., and Southwell, B.R. (2010). Cholinergic neurotransmission and muscarinic receptors in the enteric nervous system. Prog Histochem Cytochem 44, 173–202. 10.1016/j.proghi.2009.10.001.

56. Xiong, Z., Zhu, X., Geng, J., Xu, Y., Wu, R., Li, C., Fan, D., Qin, X., Du, Y., Tian, Y., et al. (2022). Intestinal Tuft-2 cells exert antimicrobial immunity via sensing bacterial metabolite N-undecanoylglycine. Immunity 55, 686–700.e7. 10.1016/j.immuni.2022.03.001.

57. Keshavarz, M., Faraj Tabrizi, S., Ruppert, A.-L., Pfeil, U., Schreiber, Y., Klein, J., Brandenburger, I., Lochnit, G., Bhushan, S., Perniss, A., et al. (2022). Cysteinyl leukotrienes and acetylcholine are biliary tuft cell cotransmitters. Sci Immunol 7, eabf6734. 10.1126/sciimmunol.abf6734.

58. Lee, R.J., Kofonow, J.M., Rosen, P.L., Siebert, A.P., Chen, B., Doghramji, L., Xiong, G., Adappa, N.D., Palmer, J.N., Kennedy, D.W., et al. (2014). Bitter and sweet taste receptors regulate human upper respiratory innate immunity. J. Clin. Invest. 124, 1393–1405. 10.1172/JCI72094.

59. Zhu, H., Aryal, D.K., Olsen, R.H.J., Urban, D.J., Swearingen, A., Forbes, S., Roth, B.L., and Hochgeschwender, U. (2016). Cre-dependent DREADD (Designer Receptors Exclusively Activated by Designer Drugs) mice. Genesis 54, 439–446. 10.1002/dvg.22949.

60. Chen, X., Choo, H., Huang, X.-P., Yang, X., Stone, O., Roth, B.L., and Jin, J. (2015). The first structure-activity relationship studies for designer receptors exclusively activated by designer drugs. ACS Chem Neurosci 6, 476–484. 10.1021/cn500325v.

61. Barilli, A., Aldegheri, L., Bianchi, F., Brault, L., Brodbeck, D., Castelletti, L., Feriani, A., Lingard, I., Myers, R., Nola, S., et al. (2021). From High-Throughput Screening to Target Validation: Benzo[d]isothiazoles as Potent and Selective Agonists of Human Transient Receptor Potential Cation Channel Subfamily M Member 5 Possessing In Vivo Gastrointestinal Prokinetic Activity in Rodents. J Med Chem 64, 5931–5955. 10.1021/acs.jmedchem.1c00065.

62. Virginio, C., Aldegheri, L., Nola, S., Brodbeck, D., Brault, L., Raveglia, L.F., Barilli, A., Sabat, M., and Myers, R. (2022). Identification of positive modulators of TRPM5 channel from a high-throughput screen using a fluorescent membrane potential assay. SLAS Discov 27, 55–64. 10.1016/j.slasd.2021.10.004.

63. Wyatt, K.D., Sakamoto, K., and Watford, W.T. (2022). Tamoxifen administration induces histopathologic changes within the lungs of Cre-recombinase-negative mice: A case report. Lab Anim 56, 297–303. 10.1177/00236772211042968.

64. Chu, C., Parkhurst, C.N., Zhang, W., Zhou, L., Yano, H., Arifuzzaman, M., and Artis, D. (2021). The ChAT-acetylcholine pathway promotes group 2 innate lymphoid cell responses and anti-helminth immunity. Sci Immunol 6, eabe3218. 10.1126/sciimmunol.abe3218.

65. Roberts, L.B., Schnoeller, C., Berkachy, R., Darby, M., Pillaye, J., Oudhoff, M.J., Parmar, N., Mackowiak, C., Sedda, D., Quesniaux, V., et al. (2021). Acetylcholine production by group 2 innate lymphoid cells promotes mucosal immunity to helminths. Sci Immunol 6, eabd0359. 10.1126/sciimmunol.abd0359.

66. C, S., Dj, T., and Rk, G. (2018). A sticky end for gastrointestinal helminths; the role of the mucus barrier. Parasite immunology 40. 10.1111/pim.12517.

67. Middelhoff, M., Nienhüser, H., Valenti, G., Maurer, H.C., Hayakawa, Y., Takahashi, R., Kim, W., Jiang, Z., Malagola, E., Cuti, K., et al. (2020). Prox1-positive cells monitor and sustain the murine intestinal epithelial cholinergic niche. Nat Commun 11, 111. 10.1038/s41467-019-13850-7.

68. Takahashi, T., Shiraishi, A., Murata, J., Matsubara, S., Nakaoka, S., Kirimoto, S., and Osawa, M. (2021). Muscarinic receptor M3 contributes to intestinal stem cell maintenance via EphB/ephrin-B signaling. Life Sci Alliance 4, e202000962. 10.26508/lsa.202000962.

69. Madden, K.B., Yeung, K.A., Zhao, A., Gause, W.C., Finkelman, F.D., Katona, I.M., Urban, J.F., and Shea-Donohue, T. (2004). Enteric nematodes induce stereotypic STAT6-dependent alterations in intestinal epithelial cell function. J Immunol 172, 5616–5621. 10.4049/jimmunol.172.9.5616.

70. Schütz, B., Jurastow, I., Bader, S., Ringer, C., von Engelhardt, J., Chubanov, V., Gudermann, T., Diener, M., Kummer, W., Krasteva-Christ, G., et al. (2015). Chemical coding and chemosensory properties of cholinergic brush cells in the mouse gastrointestinal and biliary tract. Front Physiol 6, 87. 10.3389/fphys.2015.00087.

71. Jk, G., A, E., D, A., Me, J., He, N., K, T., H, H., H, S., and Gc, H. (2012). Bicarbonate and functional CFTR channel are required for proper mucin secretion and link cystic fibrosis with its mucus phenotype. The Journal of experimental medicine 209. 10.1084/jem.20120562.

72. Birchenough, G.M.H., Johansson, M.E.V., Gustafsson, J.K., Bergström, J.H., and Hansson, G.C. (2015). New developments in goblet cell mucus secretion and function. Mucosal Immunol 8, 712–719. 10.1038/mi.2015.32.

73. Knoop, K.A., McDonald, K.G., McCrate, S., McDole, J.R., and Newberry, R.D. (2015). Microbial sensing by goblet cells controls immune surveillance of luminal antigens in the colon. Mucosal Immunol 8, 198–210. 10.1038/mi.2014.58.

74. Gustafsson, J.K., Davis, J.E., Rappai, T., McDonald, K.G., Kulkarni, D.H., Knoop, K.A., Hogan, S.P., Fitzpatrick, J.A., Lencer, W.I., and Newberry, R.D. (2021). Intestinal goblet cells sample and deliver lumenal antigens by regulated endocytic uptake and transcytosis. Elife 10, e67292. 10.7554/eLife.67292.

75. Dolan, B., Ermund, A., Martinez-Abad, B., Johansson, M.E.V., and Hansson, G.C. (2022). Clearance of small intestinal crypts involves goblet cell mucus secretion by intracellular granule rupture and enterocyte ion transport. Sci Signal 15, eabl5848. 10.1126/scisignal.abl5848.

76. Birchenough, G.M.H., Nyström, E.E.L., Johansson, M.E.V., and Hansson, G.C. (2016). A sentinel goblet cell guards the colonic crypt by triggering Nlrp6-dependent Muc2 secretion. Science 352, 1535–1542. 10.1126/science.aaf7419.

77. Morroni, M., Cangiotti, A.M., and Cinti, S. (2007). Brush cells in the human duodenojejunal junction: an ultrastructural study. J Anat 211, 125–131. 10.1111/j.1469-7580.2007.00738.x.

78. Cheng, X., Voss, U., and Ekblad, E. (2018). Tuft cells: Distribution and connections with nerves and endocrine cells in mouse intestine. Exp Cell Res 369, 105–111. 10.1016/j.yexcr.2018.05.011.

79. Hollenhorst, M.I., Kumar, P., Zimmer, M., Salah, A., Maxeiner, S., Elhawy, M.I., Evers, S.B., Flockerzi, V., Gudermann, T., Chubanov, V., et al. (2022). Taste Receptor Activation in Tracheal Brush Cells by Denatonium Modulates ENaC Channels via Ca2+, cAMP and ACh. Cells 11, 2411. 10.3390/cells11152411.

80. Banerjee, A., Herring, C.A., Chen, B., Kim, H., Simmons, A.J., Southard-Smith, A.N., Allaman, M.M., White, J.R., Macedonia, M.C., Mckinley, E.T., et al. (2020). Succinate Produced by Intestinal Microbes Promotes Specification of Tuft Cells to Suppress Ileal Inflammation. Gastroenterology 159, 2101–2115.e5. 10.1053/j.gastro.2020.08.029.

81. Huh, W.J., Roland, J.T., Asai, M., and Kaji, I. (2020). Distribution of duodenal tuft cells is altered in pediatric patients with acute and chronic enteropathy. Biomedical Research 41, 113–118. 10.2220/biomedres.41.113.

82. Aigbologa, J., Connolly, M., Buckley, J.M., and O’Malley, D. (2020). Mucosal Tuft Cell Density Is Increased in Diarrhea-Predominant Irritable Bowel Syndrome Colonic Biopsies. Front Psychiatry 11, 436. 10.3389/fpsyt.2020.00436.

83. 83. Fung, C., Fraser, L.M., Barrón, G.M., Gologorsky, M.B., Atkinson, S.N., Gerrick, E.R., Hayward, M., Ziegelbauer, J., Li, J.A., Nico, K.F., et al. (2022). Tuft cells mediate commensal remodeling of the small intestinal antimicrobial landscape. 2022.10.24.512770. 10.1101/2022.10.24.512770.

84. Sa, R., C, L., Am, M., K, L., Kl, T., C, B., S, K., Sk, G., A, N., Jm, A.-G., et al. (2019). CFTR-PTEN-dependent mitochondrial metabolic dysfunction promotes Pseudomonas aeruginosa airway infection. Science translational medicine 11. 10.1126/scitranslmed.aav4634.

85. Voehringer, D., Reese, T.A., Huang, X., Shinkai, K., and Locksley, R.M. (2006). Type 2 immunity is controlled by IL-4/IL-13 expression in hematopoietic non-eosinophil cells of the innate immune system. J Exp Med 203, 1435–1446. 10.1084/jem.20052448.

86. Me, K., Cm, M., Jg, G., Sd, P., An, G., R, A., S, Y., Pe, B., Ca, S., Pl, Z., et al. (2022). IL-13-programmed airway tuft cells produce PGE2, which promotes CFTR-dependent mucociliary function. JCI insight 7. 10.1172/jci.insight.159832.

87. Nadjsombati, M.S., McGinty, J.W., Lyons-Cohen, M.R., Jaffe, J.B., DiPeso, L., Schneider, C., Miller, C.N., Pollack, J.L., Nagana Gowda, G.A., Fontana, M.F., et al. (2018). Detection of Succinate by Intestinal Tuft Cells Triggers a Type 2 Innate Immune Circuit. Immunity 49, 33–41.e7. 10.1016/j.immuni.2018.06.016.

88. Chudnovskiy, A., Mortha, A., Kana, V., Kennard, A., Ramirez, J.D., Rahman, A., Remark, R., Mogno, I., Ng, R., Gnjatic, S., et al. (2016). Host-Protozoan Interactions Protect from Mucosal Infections through Activation of the Inflammasome. Cell 167, 444–456.e14. 10.1016/j.cell.2016.08.076.

89. Law, C.W., Chen, Y., Shi, W., and Smyth, G.K. (2014). voom: Precision weights unlock linear model analysis tools for RNA-seq read counts. Genome Biol 15, R29. 10.1186/gb-2014-15-2-r29.

90. Robinson, M.D., and Oshlack, A. (2010). A scaling normalization method for differential expression analysis of RNA-seq data. Genome Biol 11, R25. 10.1186/gb-2010-11-3-r25.

91. Robinson, M.D., McCarthy, D.J., and Smyth, G.K. (2010). edgeR: a Bioconductor package for differential expression analysis of digital gene expression data. Bioinformatics 26, 139–140. 10.1093/bioinformatics/btp616.

92. Ritchie, M.E., Phipson, B., Wu, D., Hu, Y., Law, C.W., Shi, W., and Smyth, G.K. (2015). limma powers differential expression analyses for RNA-sequencing and microarray studies. Nucleic Acids Res 43, e47. 10.1093/nar/gkv007.

